# Npas4-mediated dopaminergic regulation of fear memory states

**DOI:** 10.1101/2022.08.11.503591

**Authors:** BumJin Ko, Jong-Yeon Yoo, Woochul Choi, Rumeysa Dogan, Kibong Sung, Sangjun Lee, Dahun Um, Su Been Lee, Taesik Yoo, Hyun Jin Kim, Seung Tae Beak, Sang Ki Park, Se-Bum Paik, Tae-Kyung Kim, Joung-Hun Kim

## Abstract

Amygdala circuitry encodes associations between conditioned stimuli and aversive unconditioned stimuli, and also controls fear expression (Pape and Pare, 2010). However, whether and how irrelevant information for unpaired conditioned stimuli (CS^-^) is discretely processed, and how it was influenced by stress remain unknown. CS^-^ memory is retrievable immediately after fear conditioning, but then becomes silent after memory consolidation in mice. Synaptic pathway from the lateral to the anterior basal amygdala gates the expression of CS^-^ memory, depending upon Npas4-mediated Drd4 synthesis. The upregulation of Npas4-Drd4 axis, which is precluded by corticosterone, shifts functional states of neural engrams for CS^-^ memory toward silent states and limits its retrievability. In here, we demonstrate the cellular and molecular mechanisms regulating the functional states of neural engrams, which can be switched or maintained, supporting discriminative memory.

## Introduction

Memory operation enables organisms to adapt behaviors through the experienced information of external and internal stimuli. Among various external stimuli, the relevant information is selectively retained and represented in brains, but irrelevant information cannot be retrieved later (Terada et al., 2021). The successful dissociation between two types of information involves the reorganization of memory structures (Stickgold and Walker, 2013; Klinzing et al., 2019; Terada et al., 2021). Failure of this reorganization during threat learning often results in maladaptation of fear memory and thereby risk of survival (Holt et al., 2014). The reorganization of memory structures can be explored using discriminative Pavlovian conditioning (Ciocchi et al., 2010; Tovote et al., 2015). In this behavioral paradigm, animals are trained to distinguish cues paired with aversive stimuli such as electric shocks (CS^+^, conditioned stimulus) from ones that are not paired (CS^-^, unpaired conditioned stimulus) (Pearce, 1994). After fear conditioning, the subject animals show elevation of freezing levels toward CS^+^ but generally neglect CS^-^. The basolateral amygdala (BLA) is a critical brain region for the processing of fear memory to both CS^+^ and CS^-^ (Fendt and Fanselow, 1999). Information for CS^+^ and CS^-^ memories seemed to be processed within BLA circuits via distinct mechanisms: the BLA displayed a difference in oscillation patterns upon CS^+^ or CS^-^ presentation after fear conditioning, pointing to the possibility that CS^+^ and CS^-^ memories could be independently encoded and stored (Stujenske et al., 2014; Karalis et al., 2016). Despite the functional and behavioral importance, how CS^-^ information is processed throughout the memory operation and can be distinguished from CS^+^ at the circuit levels remains elusive.

*De novo* protein synthesis by multiple waves of gene expression is required for memory consolidation (Schafe et al, 2000; Duvarci et al, 2008). Interestingly, immediate early genes (IEGs) are a class of essential genes that show transient and rapid induction by extracellular stimuli without the necessity of protein synthesis (Yap and Greenberg, 2018). Because they are expressed instantaneously by cellular activity, IEGs including c-Fos have been extensively used to label neurons that were activated during memory formation, which are called “neural engrams” (Reijmers et al., 2007; Tonegawa et al., 2015). Besides c-Fos, neuronal PAS domain protein 4 (Npas4) is another neuron-specific IEG that could be induced exclusively by neuronal activity (Sun and Lin, 2016). Although both Npas4 and c-Fos are induced in the activated neurons, Npas4 can orchestrate synapse formation and synaptic plasticity (Lin et al., 2008; Weng et al., 2018), differently from c-Fos (Fleischmann et al., 2003; Yap et al., 2021). Importantly, it was demonstrated that non-overlapping neuronal engrams marked by either Npas4 or c-Fos were formed after fear conditioning, each of which mediates different outcomes to the expression of fear memory (Sun et al., 2020). While functional roles of Npas4 have been identified in multiple brain regions (Ramamoorthi et al., 2011; Taniguchi et al., 2017; Sun et al., 2020), its role in memory operation and the involved downstream targets in the amygdala remain largely unknown.

The reactivation of engram neurons is necessary and sufficient to evoke memory recall (Liu et al., 2012; Denny et al., 2014; for review see, Tonegawa et al., 2015). Fear generalization to novel tones with similar frequency arose in parallel with the recruitment of the same neurons that were normally activated upon exposure to CS^+^ (Grosso et al., 2018). Consistent with these findings, fear engram neurons showed input-specific synaptic plasticity in the CS^+^-responsive pathway, which contributed to an increase in the expression of fear memory (Kim and Cho, 2017). However, the mechanisms whereby engram reactivation could be restricted to specific stimuli but prevent from eliciting memory generalization have not been fully established. While reactivation of engram neurons is likely to be controlled by synaptic plasticity (Kim and Cho, 2017; Roy et al., 2017), the mechanisms underlying dissociation between CS^-^ and CS^+^ is still lacking. Since fear memory-related CS^-^ and CS^+^ are encoded within largely distinct populations (engrams) in the amygdala (Collins and Paré, 2000; Grewe et al., 2017), individual neural engrams would be synaptically or intrinsically modulated to elicit or preclude the expression of fear memory. Indeed, pharmacologic deletion of novel tone-responsive neurons in the lateral amygdala (LA) led to impairment in fear discrimination without affecting anxiety level and processing of fear extinction (Grosso et al., 2018). Hence, the cellular and molecular regulation of engram neurons for CS^+^ and CS^-^ memories could account for efficient and discriminative processing of memory information (Chelaru and Dragoi, 2008) and potentially pathological behavioral consequences that patients with fear-related diseases often exhibit even in routine environments.

Dopamine signaling can intimately control synaptic plasticity in amygdala circuits and the magnitude of fear expression (Bissière et al., 2003; Fadok et al., 2009). We previously demonstrated that dopamine receptor D4 (Drd4) in the dorsal intercalated cell mass (ITCd) of the amygdala could demarcate fear expression by regulating the inhibitory synaptic plasticity, which reduced freezing levels to less-salient experience (Kwon et al., 2015). The ablation of Drd4 resulted in excessive fear expression and generalization which are core symptoms of PTSD (Pole, 2007; Kwon et al., 2015). Interestingly, Npas4-lacking mice displayed impairment in synaptic plasticity as well as exaggerated startle responses which is an indicator of PTSD (Shalev et al., 2000; Coutellier et al., 2012; Weng et al., 2018). If Npas4 also plays certain a role in synaptic plasticity of amygdala circuits and discriminative fear memory as shown in the hippocampus and the dentate gyrus (Weng et al., 2018; Sun et al., 2020), Npas4 could play physiological and pathological roles potentially through direct/indirect interaction with Drd4. Here, we show that fear expression after exposure to CS^-^ is governed by synaptic plasticity in the pathway from the LA to the anterior basal amygdala (aBA), which depends on Npas4-mediated Drd4 synthesis, by shifting functional states of CS^-^ engrams from active to silent. The present study reveals the dissociative processing of irrelevant CS^-^ stimuli for memory operation and further provides a new molecular perspective for adaptive control of memory engrams.

## Results

### CORT modulation of CS^-^ memory retrieval

Acute exposure to stresses was shown to facilitate subsequent expression of contextual fear memory, but not auditory fear memory (Cordero et al., 2003). We found that restraint stress prior to discriminative fear conditioning (FC) could promote freezing behavior to CS^-^ without affecting CS^+^-induced fear responses (Figure 1A, 1B and Figure 1-figure supplement 1A). As expected, the serum levels of corticosterone (CORT), a major stress hormone, increased after fear conditioning in stressed mice compared to the control group (Figure 1C). To examine whether CORT genuinely elevated freezing levels to CS^-^, we administered a glucocorticoid receptor antagonist mifepristone (10 mg/kg) immediately after fear conditioning. Stressed mice that also received mifepristone exhibited lower levels of freezing to CS^-^ than vehicle-treated stressed group while those to CS^+^ were indistinguishable during training and test sessions between groups (Figure 1D, Figure 1-figure supplement 1B and 1C).

**Figure 1.**
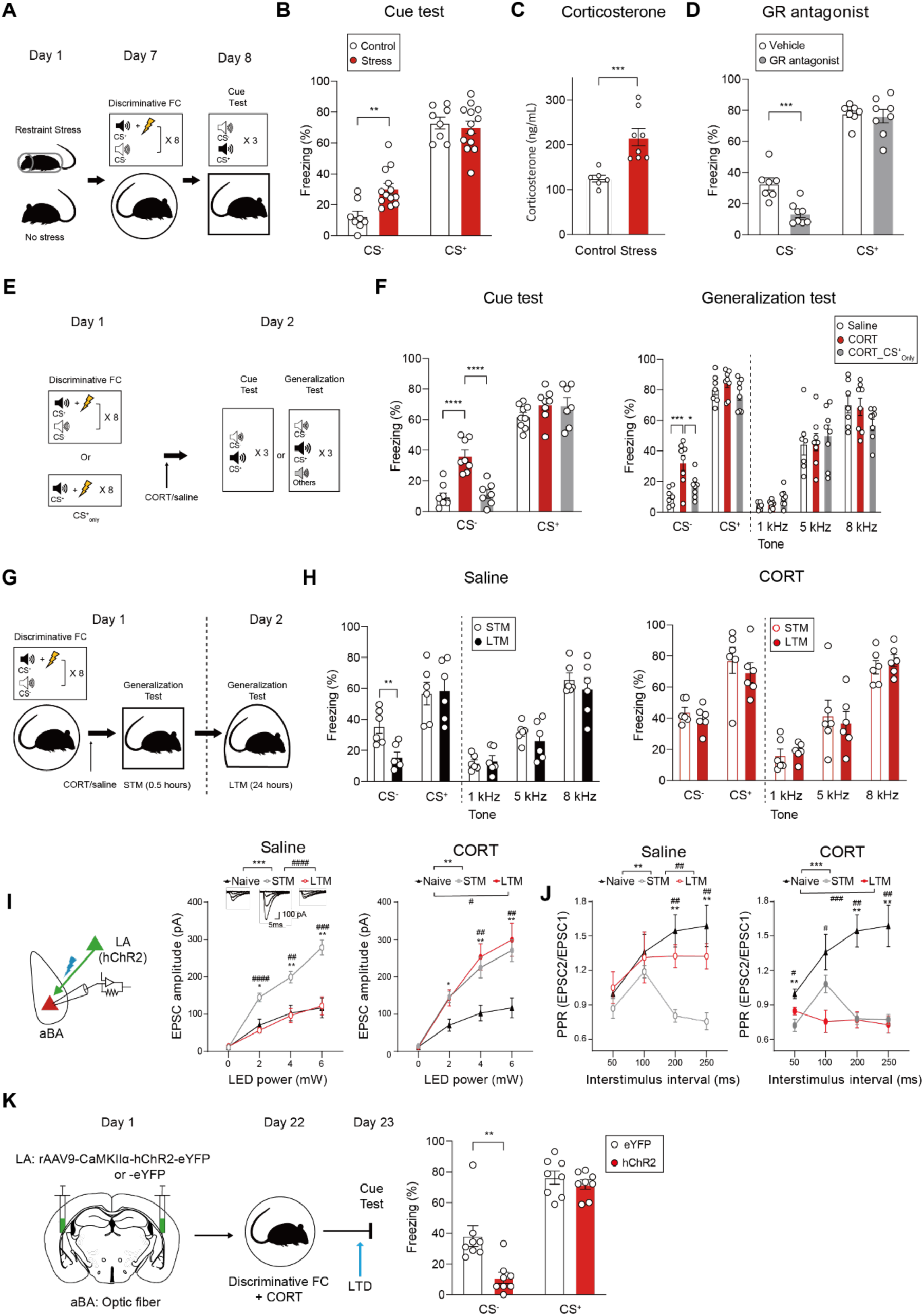
Retrievability changes of CS^-^ memory concurrent with presynaptic plasticity in the LA-to-aBA pathway. **(A)** Schematic depicting stress exposure and discriminative fear conditioning (FC). **(B)** Effects of stress exposure on fear retrieval to CS^-^ or CS^+^ (Control, n = 8; Stress, n =13 mice; Welch’s *t*-test, \*\**P* = 0.0015). **(C)** Comparison of serum CORT levels after discriminative FC between restraint stress-exposed and unexposed control mice (Control, n = 6; Stress, n = 8 mice; Mann-Whitney, \*\*\**P* = 0.0003). **(D)** Effects of GR blockage on fear responses that stressed animals displayed to presentation of CS^-^ or CS^+^ (Vehicle, n = 7; GR antagonist, n = 8 mice; Welch’s *t*-test, \*\**P* = 0.0015). **(E)** Schematic depicting fear retrieval between CORT-and saline-injected groups. **(F)** Comparison of fear responses in the cue test (left, Saline, n = 9; CORT, n = 8; CORT_CS^+^_only_, n =7; One-way ANOVA with Tukey’s *posthoc* test, \*\*\*\**P* < 0.0001) and the generalization test (right, Saline, n = 7; CORT, n = 8; CORT_CS^+^_only_, n =8; One-way ANOVA with Tukey’s *posthoc* test, \**P* = 0.0132, \*\*\**P* = 0.0009) from mice that underwent either discriminative FC or CS^+^_only_ training and then received CORT. **(G)** Schematic for generalization test at two different time points after discriminative FC. **(H)** Comparison of freezing levels to various tones at the STM and LTM time points in saline-injected (left, n = 6 mice; Paired *t*-test, \*\**P* = 0.002) and CORT-injected groups (right, n = 6 mice). **(I)** Schematic depicting *ex vivo* recording from the LA-to-aBA pathway (left). Input-output curves of optically-evoked EPSCs in saline-injected mice (middle, Naive, n = 8; STM, n = 10; LTM, n = 17 cells; Two-way repeated measures (RM) ANOVA with Sidak’s *posthoc* test, F = 24.50, \*\*\**P* = 0.0001; F = 24.44, ^####^*P* < 0.0001). Insets: representative EPSC traces at the designated time points and conditions. Input-output curves of EPSCs in CORT-injected mice (right, Naive, n = 9; STM, n = 11, LTM, n = 16 cells; Two-way RM ANOVA with Sidak’s *posthoc* test, F = 7.521, *^#^P* = 0.0119; F = 13.13, \*\**P =* 0.0021). The same data obtained from naive group were used for equal comparison. **(J)** Paired pulse ratios (PPRs) of EPSCs in saline-injected mice (left, Naive, n = 9; STM, n =10; LTM, n =10 cells; Two-way RM ANOVA followed by Sidak’s *posthoc* test, F = 14.15, \*\**P* = 0.0016; F = 11.99, ^##^*P* = 0.0028) and CORT-injected mice (right, Naive, n = 9; STM, n = 12; LTM, n = 10 cells, Two-way RM ANOVA with Sidak’s *posthoc* test, F **=** 20.03, \*\*\**P* = 0.0003; F = 18.89, *^###^P* = 0.0004). The same data obtained from naive group were used for equal comparison. **(K)** Schematic for *in vivo* induction of LTD in the LA-to-aBA pathway and subsequent behavioral test (left). Effect of LTD induction on fear responses was examined (right, eYFP, n = 8; hChR2, n = 8 mice; Welch’s *t*-test, \*\**P* = 0.0051). Data are shown as mean ± SEM.

To test whether fear responses could be directly controlled by CORT, we intraperitoneally injected either CORT (5 mg/kg) or saline to subject mice immediately after fear conditioning. CORT-injected mice froze more in response to CS^-^ than saline-injected animals while CORT injection did not affect freezing levels to CS^+^ (Figure 1E and 1F), in agreement with what we had observed following stress exposure. To ascertain whether the behavioral effects of CORT were specific to the presentation of CS^-^, we assessed fear retrieval upon exposure to CS^-^, CS^+^ or novel tones that had not been experienced. CORT-injected mice displayed higher freezing levels to CS^-^ than saline-injected group, but not to the other tones (Figure 1F). When a fraction of mice were subjected to fear conditioning without presentation of the CS^-^ and followed by CORT infusion (CORT_CS^+^_only_ group), they exhibited freezing levels comparable to those in the saline-injected group across all the tones tested (Figure 1F). CORT-injected mice showed a positive correlation of their freezing profiles between CS^+^ and 8 kHz tones but not between CS^+^ and CS^-^, which suggested the processing of CS^-^ information different from that of CS^+^ (Figure 1-figure supplement 1D and 1E). To exclude a possible bias derived from specific tone frequencies, we switched the frequency for CS^-^ and CS^+^. This counterbalanced experiment confirmed the selectivity of CORT effectiveness for the increased response to CS^-^, but not to other tones (Figure 1-figure supplement 1F). In addition, we validated the specificity of CORT effect with the lower amplitude of electric shock (low shock paradigm, 0.2 mA). First, we compared the freezing level of CS+ between low shock paradigm and our previous condition (0.4 mA). In the lower shock paradigm (0.2 mA), the mice showed lower level of freezing to CS^+^ than our previous condition (0.2 mA, n = 8 mice, 48.35 ± 3.14 %; 0.4 mA, n = 9 mice, 61.15 ± 2.32 %; Welch’s *t*-test, \*\**P* = 0.0064). Even with the lower electric shock (0.2 mA), CORT-injected mice showed higher freezing level to CS^-^, but comparable freezing level to CS^+^ to saline-injected mice (Figure 1-figure supplement 1G). Thus, the elevation of CORT during memory consolidation seemed to promote fear expression selectively to the CS^-^ stimulus experienced by subject animals during training, rather than causing generalized fear expression to a range of cues.

Neuronal ensembles encoding memories, also called neural engrams, have been postulated to have several states in accordance with their accessibility: encoded memory could be readily retrieved when their engrams are in active states but are no longer retrieved by natural stimulation when the corresponding engrams are in unavailable or silent states (Josselyn and Tonegawa, 2020). Interestingly, neural engrams in silent states, which could be recalled by artificial activation, were observed in a wide range of circumstances such as retrograde amnesia and systems consolidation (Roy et al., 2017; Kitamura et al., 2017). We reasoned that the retrievability of CS^-^ memory could be modulated or defined by individual functional states of the engram neurons. If that is the case, the observed increasement of fear responses to CS^-^ would result from CORT-induced interference in the conversion of engram states from active to silent, leading to the sustained retrievability of CS^-^ memory. To address this notion, we examined whether CS^-^ memory underwent a state transition before and after memory consolidation at the time points corresponding to STM (short-term memory, 0.5 hours after fear conditioning) and LTM (long-term memory, 24 hours). Saline-injected animals displayed decremental freezing levels to CS^-^ from the STM to LTM time points whereas freezing levels to other tones including CS^+^ remained unaltered (Figure 1G and 1H). In addition, CORT_CS^+^_only_ group showed lower level of CS^-^ freezing at both the STM and LTM time points (Figure 1-figure supplement 1H). The result indicates CORT-injection alone could not induce generalized fear in neither of time points. To test the ability of discrimination between CS^-^ and CS^+^, we compared the discrimination index. The groups showing higher level of freezing to CS^-^ also showed significantly lower level of the discrimination index compared to controls (Figure 1-figure supplement 2). Thus, CS^-^ memory appeared to be retrievable prior to memory consolidation but became inaccessible afterward. The sustained fear responses to CS^-^ observed in CORT-injected mice supported the possibility that CORT would keep the retrievability of CS^-^ memory elevated throughout behavioral tests.

### Fear expression to CS^-^ gated by the LA-to-aBA pathway

To identify which brain region(s) was(were) primarily involved in the processing of CS^-^ memory, we compared numbers of c-Fos-expressing cells in various brain regions after CS^-^ exposure. Consistent with a previous report that BLA neurons would also retain CS^-^-related information (Genud-Gabai et al., 2013), CORT-injected mice displayed higher numbers of c-Fos-expressing cells in the aBA compared to those in saline-injected group (Figure 1-figure supplement 3A and 3B). However, we observed no difference in the numbers of c-Fos-expressing cells between groups in the other brain regions including the lateral amygdala (LA), the central amygdala (CeA), the prelimbic cortex (PL) and the infralimbic cortex (IL) (Figure 1-figure supplement 3C). c-Fos-expressing cells were widely distributed along the anterior to posterior regions of the BA when mice were subjected to CS^+^, without any obvious difference between groups (Figure 1-figure supplement 3D-3F). Therefore, the aBA was likely to participate in the processing and retrieval of CS^-^-related information that was made accessible by CORT administration.

Given the potent innervation from the LA to the BA (Duvarci and Pare, 2014), we analyzed *ex vivo* synaptic transmission in the LA-to-aBA pathway at the time points for STM and LTM, respectively. To achieve the pathway-specific stimulation, we infused adeno-associated virus (AAV) encoding channelrhodopsin-2 (ChR2) under the CaMKII*α* promoter into the LA and then performed whole-cell recording from principal neurons of the aBA (Figure 1I and Figure 1-figure supplement 4A). Optical stimulation of LA axon terminals in the aBA reliably evoked monosynaptic excitatory postsynaptic currents (EPSCs) (Figure 1-figure supplement 4B and 4C). The mean amplitudes of EPSCs at the STM point in fear-conditioned animals were higher than those in naive mice. At the LTM time point, however, EPSC amplitudes recorded from fear-conditioned mice were comparable to those from naive animals, indicative of possible depotentiation of EPSCs from STM to LTM points (Figure 1I). Importantly, CORT-injected animals exhibited higher EPSC amplitudes than naive mice at both time points (Figure 1I). To parse the attributes of synaptic enhancement, we analyzed paired-pulse ratios (PPRs) of EPSCs. PPRs from saline-injected group were lower at the STM point than those from naive mice whereas CORT-injected animals showed lower levels of PPRs at both time points (Figure 1J). Furthermore, ratios of AMPAR/NMDAR-EPSCs (α-amino-3-hydroxy-5-methyl-4-isoxazolepropionic acid receptor/N-methyl-D-aspartate receptor-mediated EPSCs) were comparable among groups (Figure 1-figure supplement 4D), arguing the presynaptic nature of the observed synaptic plasticity. The decay time constant of NMDAR-mediated EPSCs in the presence of MK-801 was also smaller in CORT-LTM group than in saline-LTM group (Figure 1-figure supplement 4E). Importantly, stressed mice also exhibited higher amplitudes of EPSCs than the control group at the LTM time point, as in CORT-injected mice (Figure 1-figure supplement 4F). Taken together, presynaptic alteration of transmission efficacy in the LA-to-aBA pathway is concurrent with the magnitude changes and could account for the expression of CS^-^ memory, at least for STM and LTM .

Since potentiated synapses could be readily depressed (Yang and Faber, 1991), we applied a subthreshold protocol for long-term depression (LTD) onto the same pathway. LTD was induced in CORT-injected animals at the LTM time point, but neither in saline-injected nor naive mice. LTD was associated with increases in PPRs, which reflected a reduction of transmitters release (Figure 1-figure supplement 4G and 4H). Following *ex vivo* and *in vivo* validation of another LTD protocol in the absence of picrotoxin (Figure 1-figure supplement 4I and 4K), we optically induced *in vivo* LTD in the LA-to-aBA pathway of behaving animals. Supporting the behavioral consequence of synaptic transmission and plasticity, freezing levels to CS^-^ in CORT-injected mice became reduced after the induction of LTD compared to those in eYFP control animals whereas freezing levels to CS^+^ remained comparable in both groups (Figure 1K and Figure 1-figure supplement 4J). These data indicated that synaptic transmission in the LA-to-aBA pathway might selectively modulate fear responses to CS^-^: the enhanced transmission would result in sustained retrievability of CS^-^ memory to promote the long-lasting defensive responses.

### Distinct functional states for CS^-^ memory revealed by measurement of activity and neural representations of aBA^Rspo2(+)^ neurons

A majority of principal neurons of the aBA express Rspo2 (aBA^Rspo2(+)^ neurons), which critically contribute to aversive memory (Kim et al., 2016). To obtain mechanistic insights into their roles for the processing of CS^-^ or CS^+^ information, we monitored Ca^2+^ levels from individual aBA^Rspo2(+)^ neurons through a micro-endoscope during the fear retrieval session (Figure 2A, Figure 2-figure supplement 1A and 1B). aBA^Rspo2(+)^ neurons were categorized into three groups based on their activity toward stimuli: excited (E), inhibited (I), and non-responsive (NR) groups. The proportional size of each group and its spatial distribution were comparable between saline- and CORT-injected animals (Figure 2B, 2C and Figure 2-figure supplement 1C). Importantly, the aBA neurons belonging to group-E displayed higher activity upon CS^-^ presentation in CORT-injected mice than those in saline-injected group (Figure 2B). However, either the activity of group-I to CS^-^ or that of all the groups to CS^+^ was indistinguishable between groups (Figure 2B, 2C and Figure 2-figure supplement 1D). CORT infusion was able to elevate excitation magnitudes in aBA^Rspo2(+)^ neurons of group-E to the presentation of CS^-^ while it did not affect the proportion or distribution of each neuronal group. Notably, CS^-^-induced activity of group-E neurons was significantly attenuated after memory consolidation in saline-injected mice in agreement with our *ex vivo* recording data (Figure 1I, Figure 2-figure supplement 1E and 1F), but not in CORT-injected mice whereas the activity of group-E neurons to other cues including CS^+^ remained unaffected (Figure 2-figure supplement 1G-1J).

**Figure 2.**
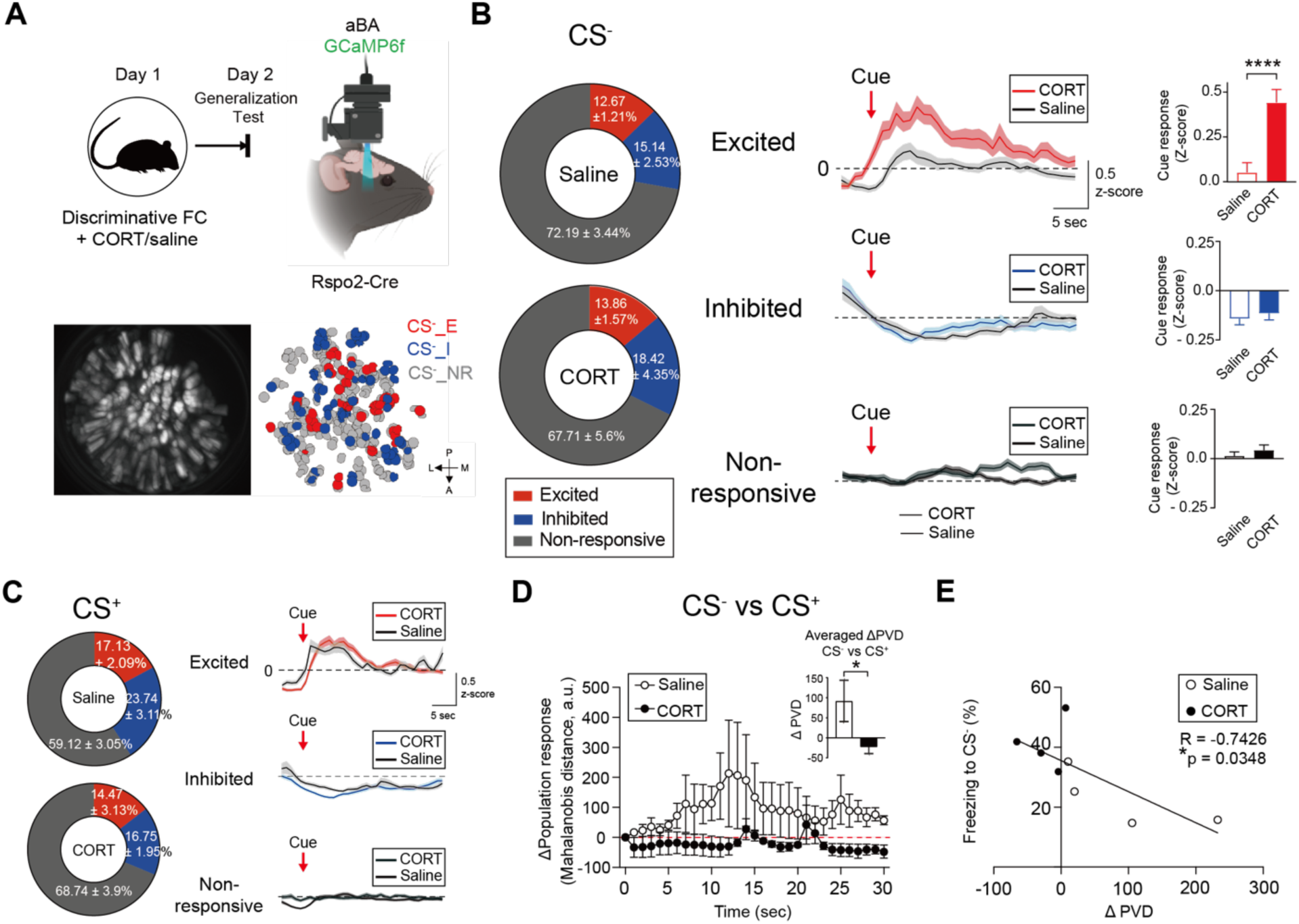
CORT-induced alteration of CS^-^-neural representation and activity. **(A)** Schematic for *in vivo* Ca^2+^ imaging from Rspo2-expressing neurons in the BA (top). A representative image of the maximum projection of field-of-view (bottom left) and extracted individual responses of Rspo2-expressing neurons toward CS^-^ (bottom right). **(B)** Classification of aBA^Rspo2(+)^ neurons in accordance with their cellular activity upon CS^-^ presentation: excited (E, red), inhibited (I, blue) and non-responsive neurons (NR, grey). The proportion of each neuronal cluster (E, I and NR to CS^-^) in saline- and CORT-injected groups (left, Saline, n = 4; CORT, n = 4 mice). Averaged z-scored activity traces of aBA^Rspo2(+)^ neurons belonging to each cluster upon CS^-^ presentation between saline- and CORT-injected mice (middle). Mean activity of each cluster to CS^-^ (right, Saline CS^-^_E, n = 91 *vs* CORT CS^-^_E, n = 98; Saline CS^-^_I, n = 105 *vs* CORT CS^-^_I, n = 112; Saline CS^-^_NR, n = 481 *vs* CORT CS^-^_NR, n = 353 cells; Welch’s *t*-test, \*\*\*\**P* < 0.0001). **(C)** The proportion of each neuronal cluster (E, I and NR to CS^+^) in saline- and CORT-injected groups (left, Saline, n = 4; CORT, n = 4 mice). Averaged z-scored activity traces of aBA^Rspo2(+)^ neurons belonging to each cluster upon CS^+^ presentation between saline- and CORT-injected mice (right). **(D)** Deviation of population vector distance (ΔPVD) between CS^-^ and CS^+^ in saline- and CORT-injected mice. Inserts: mean ΔPVD between saline- and CORT-injected mice (Saline, n = 4; CORT, n = 4 mice; Mann-Whitney test, \**P* = 0.037). **(E)** Inverse correlation between individual ΔPVD and corresponding freezing levels to CS^-^. Data are shown as mean ± SEM.

While individual neurons tend to show stochastic dynamics, neuronal population activity reliably supports fear memory possibly by encoding perceptual representations (Grewe et al., 2017). To assess the differentiability of neural responses, we computed the deviation of population vector distance (ΔPVD) between CS^-^ and CS^+^ presentations. The ΔPVD was larger in saline-injected mice than in the CORT-injected group (Figure 2D). We also measured ΔPVD for CS^+^ *vs.* each tone to ascertain specificity of the altered neural representations to exposure of CS^-^ (Figure 2-figure supplement 2A-2D) but failed to observe apparent difference regardless of CORT injection, which verified selectivity of the CS^-^-induced alteration in neural representation. If the neural representation of aBA^Rspo2(+)^ neurons to sensory cues was a critical determinant for the emergence of fear responses, ΔPVD could predict and underlie freezing behaviors upon exposure to CS^-^. Indeed, ΔPVD between CS^-^ and CS^+^ was inversely correlated with magnitudes of CS^-^-induced freezing (Figure 2E). Moreover, CS^-^-induced freezing levels could be predicted by ΔPVD measured at the STM and LTM points in saline-injected mice (Figure 2-figure supplement 2E and 2F). Therefore, our *in vivo* Ca^2+^ imaging substantiated the existence of distinct functional states in which neural ensembles for CS^-^ memory rested and further suggested that the resemblance of CS^-^ representation to CS^+^ one could demarcate the range of freezing behaviors when exposed to CS^-^.

### Upregulation of Npas4-Drd4 axis during memory consolidation

The findings for selective alternation of CS^-^-induced activity and neural representation by CORT administration prompted us to investigate molecular mechanisms underlying the synaptic and neuronal modification. As IEGs drive activity-dependent gene expression crucial for modification of synaptic transmission and plasticity (Yap and Greenberg, 2018), we began to examine which IEGs were differentially induced after fear conditioning between saline- and CORT-injected mice. When assessed at various time points, *c-Fos* and *Arc* mRNA amounts in the amygdala tissues were initially increased and subsequently resumed to baseline, which did not differ between groups (3Figure A and 3B). Importantly, the expression of *Npas4* mRNA was significantly reduced in CORT-injected mice compared to that in saline-injected group at 0.5 hours after fear conditioning, but not at later time points (Figure 3B). The time period for reduction of *Npas4* mRNA was of interest since memory consolidation occurs in a similar temporal window (David and Squire, 1984). We further examined amygdala subregions where IEGs were differentially induced by fear conditioning. Interestingly, numbers of Npas4-expressing cells were significantly smaller in the LA of CORT-injected mice than of saline-injected animals, but not in the ITCd and the BA. However, c-Fos expression in the respective amygdala areas was comparable between saline- and CORT-injected mice (Figure 3C and Figure 3-figure supplement 1A-1C).

**Figure 3.**
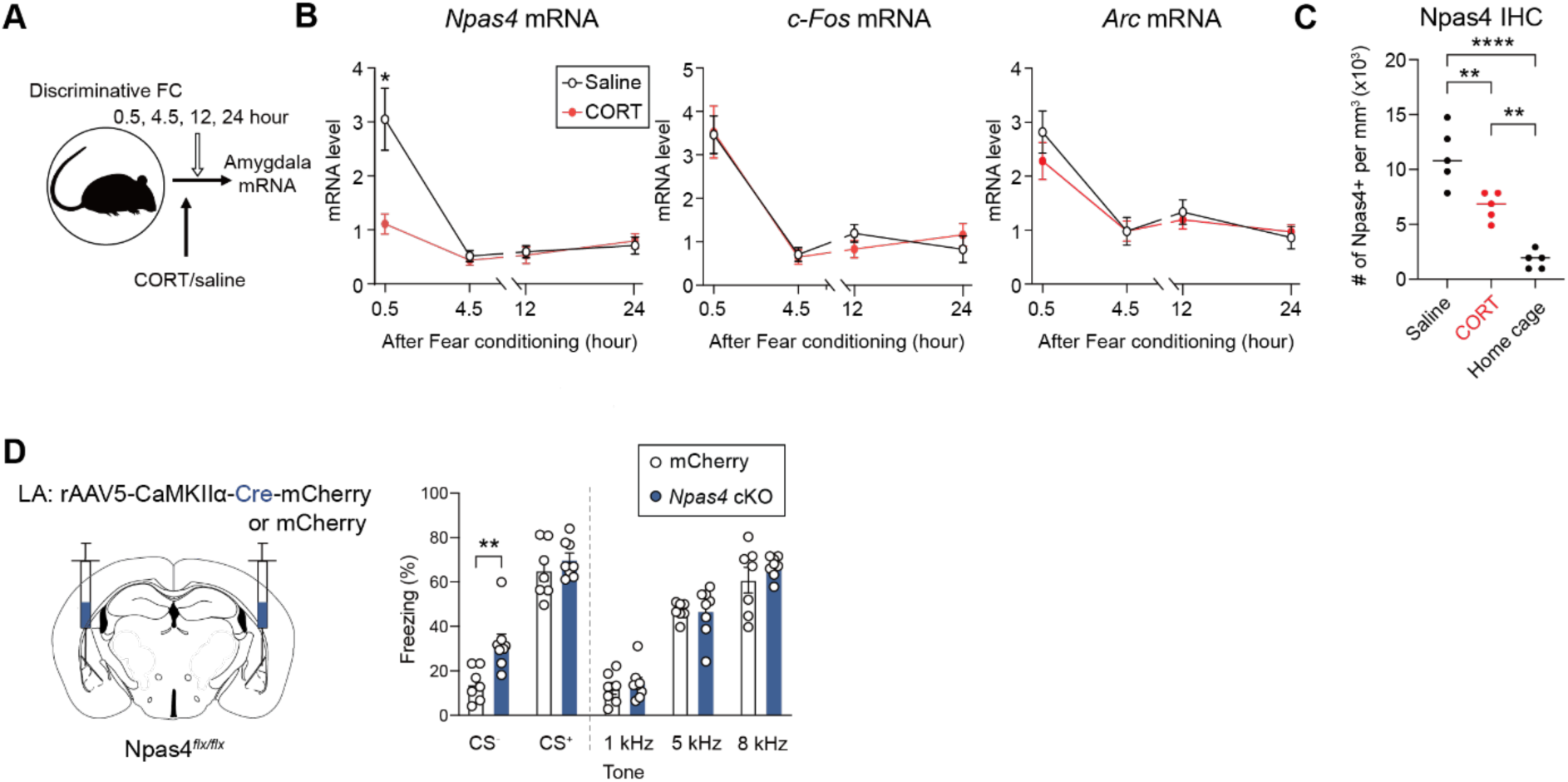
Npas4-mediated regulation of expression of CS^-^ memory. **(A)** A schematic timeline for mRNA extraction after behavioral and pharmacological interventions. **(B)** Temporal quantification of IEGs mRNAs from the amygdala tissues between saline- and CORT-injected mice after discriminative FC. mRNA amounts for individual IEGs were indicated relative to those of naive mice. The relative amounts of *Npas4* (left), *c-Fos* (middle) and *Arc* (right) mRNAs throughout time points. (Saline, n = 6; CORT, n = 6 mice; Mann-Whitney test, \**P* = 0.0303). **(C)** Numbers of Npas4-expressing cells in the LA through immunohistochemistry (IHC) (Home cage, n = 5; Saline, n = 5; CORT, n = 5 mice; One-way ANOVA with Tukey’s *posthoc* test, \*\**P* < 0.01, \*\*\*\**P* < 0.0001). **(D)** Schematic for conditional knockout (cKO) of *Npas4* in the LA principal neurons (left). Fear responses to various tones including CS^-^ and CS^+^ between *Npas4* cKO and mCherry control mice (right, mCherry, n = 7; *Npas4* cKO, n = 8 mice; Welch’s *t*-test, **P = 0.0045). Data are shown as mean ± SEM.

To elucidate the functional roles of Npas4 and further recapitulate the reduced expression of Npas4 observed in CORT-injected mice, we conditionally knocked out (cKO) *Npas4* in the LA of Npas4-floxed mice by injecting AAV encoding Cre recombinase under the CaMKIIα promoter and then assessed CS^-^-induced fear responses. *Npas4* cKO mice displayed higher freezing levels to CS^-^ than mCherry group, but no apparent alteration in freezing responses to the other tones (Figure 3D) and lower discrimination index (Figure 3-figure supplement 1D). Despite changes in CS^-^-induced responses, there was no difference in freezing profiles during fear conditioning between groups as well as anxiety or pain sensitivity (Figure 3-figure supplement 1D-1F). Interestingly, *Npas4* cKO mice showed higher amplitudes of EPSCs than mCherry group at the LTM time point (Figure 3-figure supplement 1G), as shown in CORT-injected mice. Thus, Npas4 expression would at least partly reduce or prevent the CS^-^-induced fear responses from being exaggerated likely by keeping the synaptic transmission in a normal range.

As a transcription factor, Npas4 regulates the expression of multiple genes to orchestrate activity-dependent genetic programs (Ramamoorthi et al., 2011). To elucidate downstream targets of Npas4, we performed CUT&Tag experiments (Kaya-Okur et al., 2019; see also methods for experimental details) with the amygdala tissues isolated from fear-conditioned mice (Figure 4A). We identified 54,493 of Npas4-enriched regions in which Npas4 binding motif was represented (Figure 4-figure supplement 1A-1C). Out of these, we focused on *Drd4* (dopamine receptor D4) since it could control fear responses specifically to less-salient stimuli (Kwon et al., 2015). The *Drd4* promoter region showed robust enrichment of Npas4 binding sites (Figure 4B). In addition, the luciferase assay using *Drd4* promoters revealed that their activity was significantly increased in the presence of Npas4. Deletion of both the postulated Npas4 motifs and the Npas4 binding sites (identified through CUT&Tag analysis) within the *Drd4* promoter region resulted in a decrease in luciferase activity (Figure 4C), supporting binding of Npas4 to the promoter. In fact, *Drd4* mRNA amounts increased from 4.5 hours and were kept elevated until 24 hours after fear conditioning in saline-injected mice, but not in CORT-injected group (Figure 4D). Our fluorescence *in situ* hybridization (FISH) and immunohistochemistry (IHC) corroborated that *Drd4* mRNA and Drd4 protein levels were higher in the LA and the ITCd of saline-injected mice than those of CORT-injected mice (Figure 4E, 4F and Figure 4-figure supplement 2A). The temporal expression profiles of *Npas4* and *Drd4* mRNAs (Figure 3B and 4D) were also consistent with Npas4-mediated transcriptional regulation of *Drd4*, which would be affected by CORT elevation during memory consolidation. Altogether, Npas4 could promote Drd4 transcription by direct binding to the Drd4 promoter. To assess the behavioral consequences of the cell type- and region-specific depletion of Drd4, we attempted to deplete Drd4 specifically at the LA pyramidal neurons of CaMKIIα-Cre mice by micro-infusion of AAV containing Cre-dependent small hairpin RNA (shRNA) against *Drd4* (Kwon et al., 2015) and then examined fear responses to CS^-^ (Figure 4G). As observed in *Npas4* cKO mice, depletion of Drd4 in the LA led to increases in freezing levels to CS^-^ and reduced discrimination index, without affecting fear conditioning and fear responses to other tones (Figure 4H, 4I and Figure 4-figure supplement 2B) as well as anxiety levels or pain sensitivity (Figure 4-figure supplement 2C and 2D).

**Figure 4.**
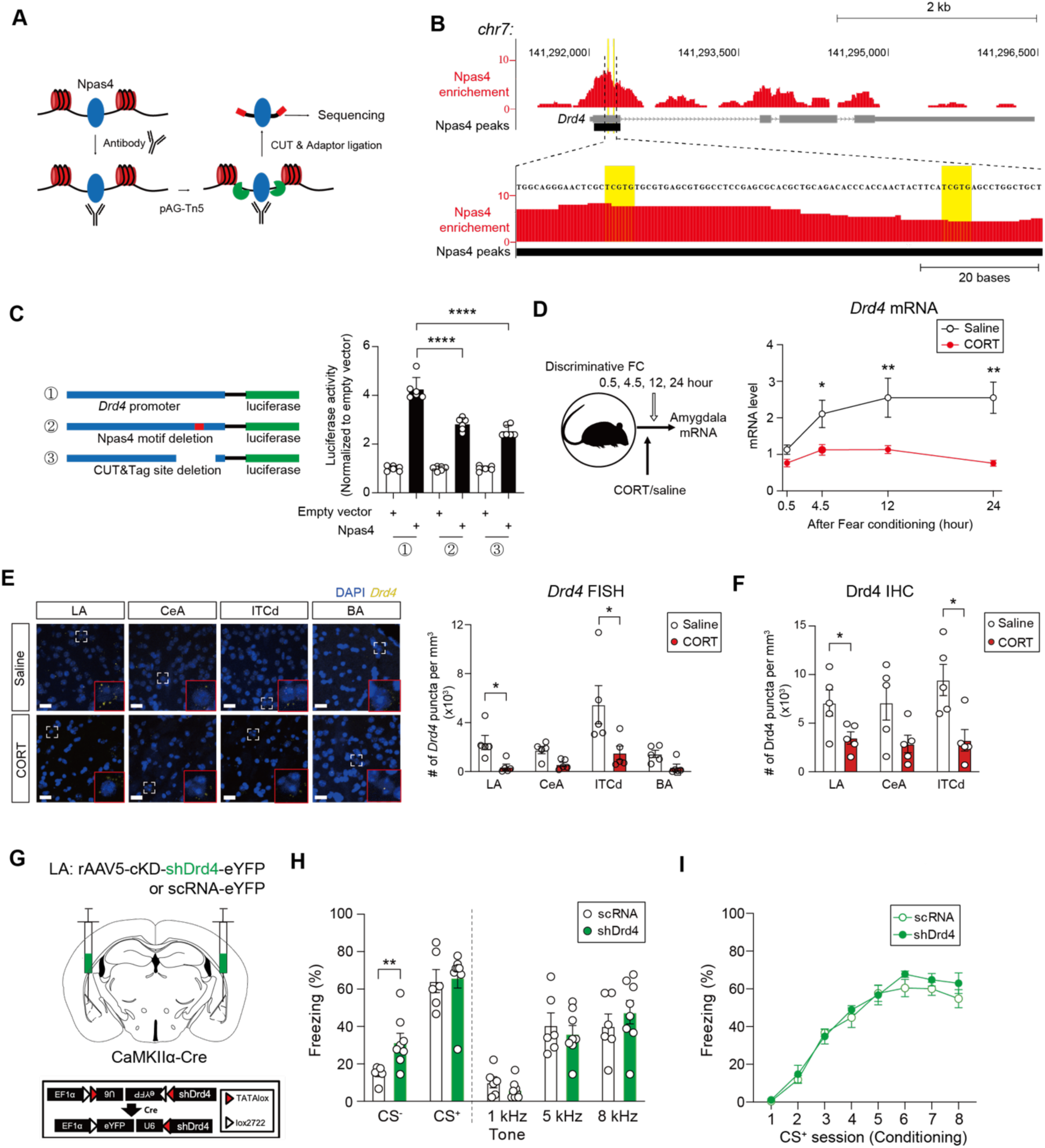
Aberrant expression of CS^-^ memory precluded by upregulation of Npas4-Drd4 axis. **(A)** A schematic diagram depicting CUT&Tag assay for Npas4 signaling. **(B)** Binding sites of Npas4 at the *Drd4* promoter region delineated by CUT&Tag (top). Npas4 motif (TCGTG) within Npas4 peaks are highlighted (yellow, bottom). Y-axis indicates normalized reads (10^7^). **(C)** Wild-type (WT) or mutant *Drd4* promoter-containing probes used for luciferase assays: ① the WT Drd4 promoter, ② the mutant promoter lacking postulated Npas4 motif and ③ the mutant promoter lacking Npas4 binding sites identified by CUT&Tag assay (left). *Drd4* promoter activity was analyzed for each condition (right) (①, n = 6; ②, n = 6; ③, n = 6 wells; One-way ANOVA with Tukey’s *posthoc* test, \*\*\*\**P* < 0.0001). **(D)** Temporal quantification of *Drd4* mRNA from the amygdala tissues obtained after discriminative FC between saline- and CORT-injected mice. A schematic timeline for mRNA collection (left). *Drd4* mRNA amounts relative to those of naive mice throughout time points (right, Saline, n = 6; CORT, n = 6 mice; Mann-Whitney test, \**P* < 0.05, \*\**P* < 0.01). **(E)** Representative images of fluorescence puncta for *Drd4* mRNA 24 hours after discriminative FC in the LA, the CeA, the ITCd and the BA of saline- and CORT-injected mice. Areas outlined with white dotted lines are magnified into those with red solid lines (left). Scale bars: 20 μm. Quantified data of *Drd4* RNA punctum density in sub-nuclei of the amygdala were compared between saline- and CORT-injected mice (right, Saline, n = 5; CORT n = 5 mice; Mann-Whitney test, \**P* < 0.05). **(F)** Drd4 punctum density measured in sub-nuclei of the amygdala 24 hours after discriminative FC (Saline, n = 5; CORT, n = 5 mice; Mann-Whitney test, \**P* < 0.05). **(G)** Schematic for Drd4 conditional knockdown (cKD) in the LA (top) and a schematic diagram for the shRNA probe used for Drd4 depletion (bottom). **(H)** Effects of Drd4 depletion in LA principal neurons on fear responses to various tones in the generalization test. Viruses containing either shRNA-targeting *Drd4* (shDrd4) or scrambled RNA (scRNA) as a negative control were micro-infused into the LA (shDrd4, n = 8; scRNA, n = 6; Welch’s *t*-test, \*\**P* = 0.0083). **(I)** Freezing profiles during FC between shDrd4- and scRNA-injected mice (shDrd4, n = 8; scRNA, n = 6 mice). Data are shown as mean ± SEM.

### Processing of CS^-^ memory gated by Drd4-mediated synaptic regulation

Based on the evidence that both Drd4 expression and synaptic transmission in the LA-to-aBA pathway were controlled by CORT and Npas4, we inferred that Drd4 signaling itself could modulate synaptic transmission. We perfused a Drd4 agonist (PD-168,077) while recording EPSCs from aBA neurons. The activation of Drd4 induced LTD only in CORT-LTM group but not in other groups (Figure 5A). However, the perfusion of Drd1 agonist (SKF-38393) did not affect synaptic transmission in CORT-LTM mice (Figure 5-figure supplement 1A). We also turned to *Drd4* KO mice to examine possible off-target effects of the used Drd4 agonist but did not observe any alteration of synaptic transmission in the same pathway despite prior injection of CORT (Figure 5-figure supplement 1B). In addition, LTD was induced by perfusion of dopamine (30 µM) in CORT-LTM group, which was totally abolished in the presence of a Drd4 antagonist (L-745,870) (Figure 5-figure supplement 1C).

**Figure 5.**
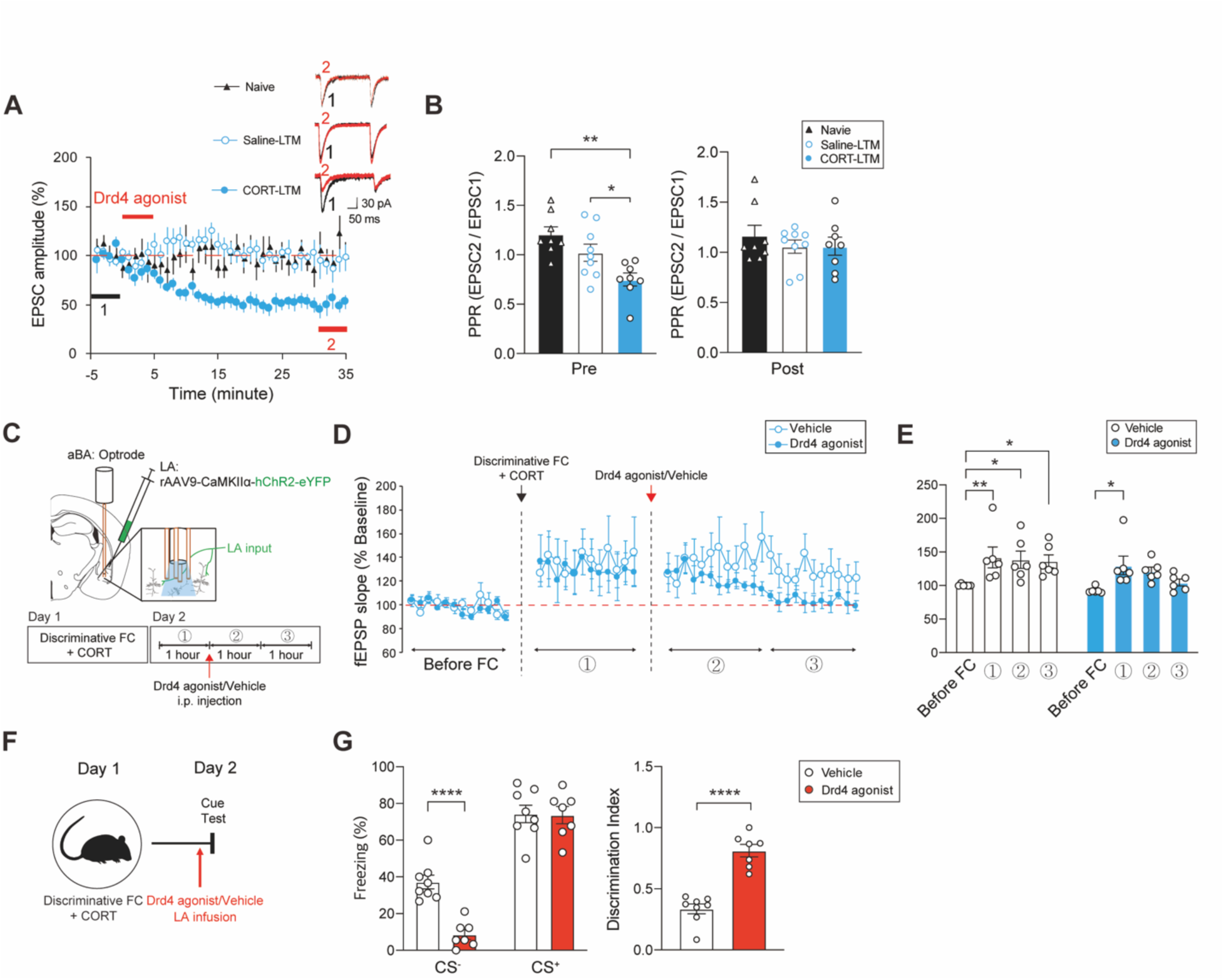
Drd4-induced depotentiation of the LA-to-aBA pathway. **(A)** Effects of Drd4 agonist (PD-168,077) on synaptic transmission of the LA-to-aBA pathway of naive, saline- or CORT-injected mice at the LTM time point. Inserts: representative traces of EPSCs at the color-designated time points (Naive, n = 8; Saline-LTM, n = 9; CORT-LTM, n = 8 cells). **(B)** PPR measurement (250 ms) of the evoked EPSCs before (left) and after PD-168,077 application (right)(Naive, n = 8; Saline-LTM, n = 9; CORT-LTM, n = 8 cells; One-way ANOVA with Tukey’s *posthoc* test, \**P* = 0.0498, \*\**P* = 0.0015). **(C)** Schematic (top) and an experimental timeline (bottom) for *in vivo* recording of fEPSPs from the aBA while stimulating LA axon terminals. **(D)** fEPSP slopes before and after discriminative FC with CORT injection were measured with/without subsequent treatment of PD-168,077 (Vehicle, n = 6; Drd4 agonist, n = 6 mice). **(E)** Averaged fEPSP slopes for each recording session (Vehicle, n = 6; Drd4 agonist, n = 6 mice; One-way repeated measures (RM) ANOVA with Tukey’s *posthoc* test, *\*P* = 0.0189). **(F)** An experimental timeline for micro-infusion of either Drd4 agonist (PD-168,077) or vehicle into the LA of animals that underwent discriminative FC and CORT injection, and behavioral tests. **(G)** Effect of PD-168,077 micro-infusion on fear retrieval in the cue test (left, Vehicle, n = 8; Drd4 agonist, n = 7 mice; Welch’s *t*-test, \*\*\*\**P* < 0.0001) and discrimination index (right, Welch’s *t*-test, \*\*\*\**P* < 0.0001). Data are shown as mean ± SEM.

We explored the synaptic attributes of Drd4-induced LTD that occurred in CORT-injected mice. Inhibition of GPCR signaling with GDPβS included in recording pipettes did not affect LTD, arguing against the possibility that LTD resulted from alteration of postsynaptic neurons (Figure 5-figure supplement 1D). Drd4-induced LTD was accompanied by increasement of PPRs in CORT-LTM group (from 0.75 ± 0.065 to 1.061 ± 0.09, mean ± SEM). PPRs measured from all the groups became similar by application of PD-168,077 (Figure 5B). Moreover, we assessed Drd4 expression in individual axon terminals of LA principal neurons within the aBA via immuno-electron microscopic (EM) imaging. To identify the very principal neurons for immuno-EM imaging, we injected AAV encoding hChR2-eYFP under the CaMKIIα promoter into the LA, which was subsequently labelled with anti-GFP antibody. Immuno-labelled Drd4 was mainly detected in the presynaptic sites of asymmetric synapses formed by the LA neurons elucidated by eYFP signals (Figure 5-figure supplement 1E).

Given the fact that synaptic transmission of the LA-to-aBA pathway was potentiated by CORT but depressed by Drd4 activity (Figure 1I and 5A), we hypothesized that the synaptic transmission could be normalized (or depotentiated) *in vivo* by Drd4 signaling. To address this notion, we recorded local field potential in behaving animals while optically stimulating LA axon terminals in the aBA (Figure 5C). Indeed, slopes of *in vivo* field EPSPs (fEPSPs) increased after fear conditioning in CORT-injected mice but resumed to the baselines after systemic treatment of PD-168,077. However, the slopes of fEPSPs in vehicle-treated mice remained elevated in the same conditions (Figure 5D and 5E). We tested whether activation of Drd4 affected synaptic transmission at the STM time point when synaptic potentiation was manifest in both saline- and CORT-injected mice. LTD could be induced in both groups (Figure 5-figure supplement 1F and 1G), which corroborated the efficacy of Drd4 activity for reduction of the increased transmission. Interestingly, we found that PD-168,077 readily induced LTD in stress-LTM group (Figure 5-figure supplement 1H) but not in saline-LTM group, indicative of occlusion of LTD. Based on the aforementioned physiological data, we began to assess whether Drd4 activation was sufficient to normalize or reduce fear retrieval to CS^-^ in CORT-injected mice. When micro-infused with PD-168,077 or vehicle into the LA (Figure 5F), mice that received PD-168,077 exhibited lower freezing levels to CS^-^ and higher discrimination index than vehicle-treated control group, without affecting fear responses to CS^+^ (Figure 5G). Collectively, LTD of the LA-to-aBA pathway seemed to be mediated mainly by activation of presynaptically-localized Drd4 during memory consolidation and in turn, contributed to the diminution of fear retrieval to CS^-^.

### Functional states of CS*^-^* memory governed by activity of Npas4-expressing neurons

Since Npas4-induced synthesis of Drd4 could lower the retrievability of CS^-^ memory (Figure 4), the activity of Npas4-expressing neurons in the LA could be sufficient to elaborate the functional shift of neural engrams for CS^-^ memory from silent states to active ones, which could result in leading to elevation of freezing levels of CS^-^ (Figure 6A). We sought to directly mark and then manipulate the activity of Npas4-expressing neurons using the *N*-RAM reporter combined with doxycycline (Dox)-dependent Tet-off system (Sun et al., 2020): tTA (tetracycline-controlled transactivator) was driven by the *N*-RAM promoter, which resulted in expression of DREADDs (hM3Dq or hM4Di) in Dox-off period (Figure 6B and Figure 6-figure supplement 1A-1E). While leak expression was negligible in Dox-on (FC + Dox) group, numbers of *N*-RAM-labelled neurons within the LA area were higher in Dox-off (FC) group than those in home cage (HC) group (Figure 6C). Importantly, the mice harboring hM3Dq displayed higher freezing levels to CS^-^ than control mice, but similar fear responses to other tones (Figure 6D), suggesting the requirement of activity of Npas4-expressing neurons in curtailing the retrievability of CS^-^ memory. However, freezing levels to all tested tones were indistinguishable in animals expressing hM4Di or mCherry in the Npas4-expressing neurons of the LA (Figure 6D) presumably due to their intrinsically low activity, which also indicated that they were dispensable for fear expression to CS^+^. Also, the hM3Dq group showed lower level of discrimination index to both mCherry and hM4Di group (Figure 6-figure supplement 1C).

**Figure 6.**
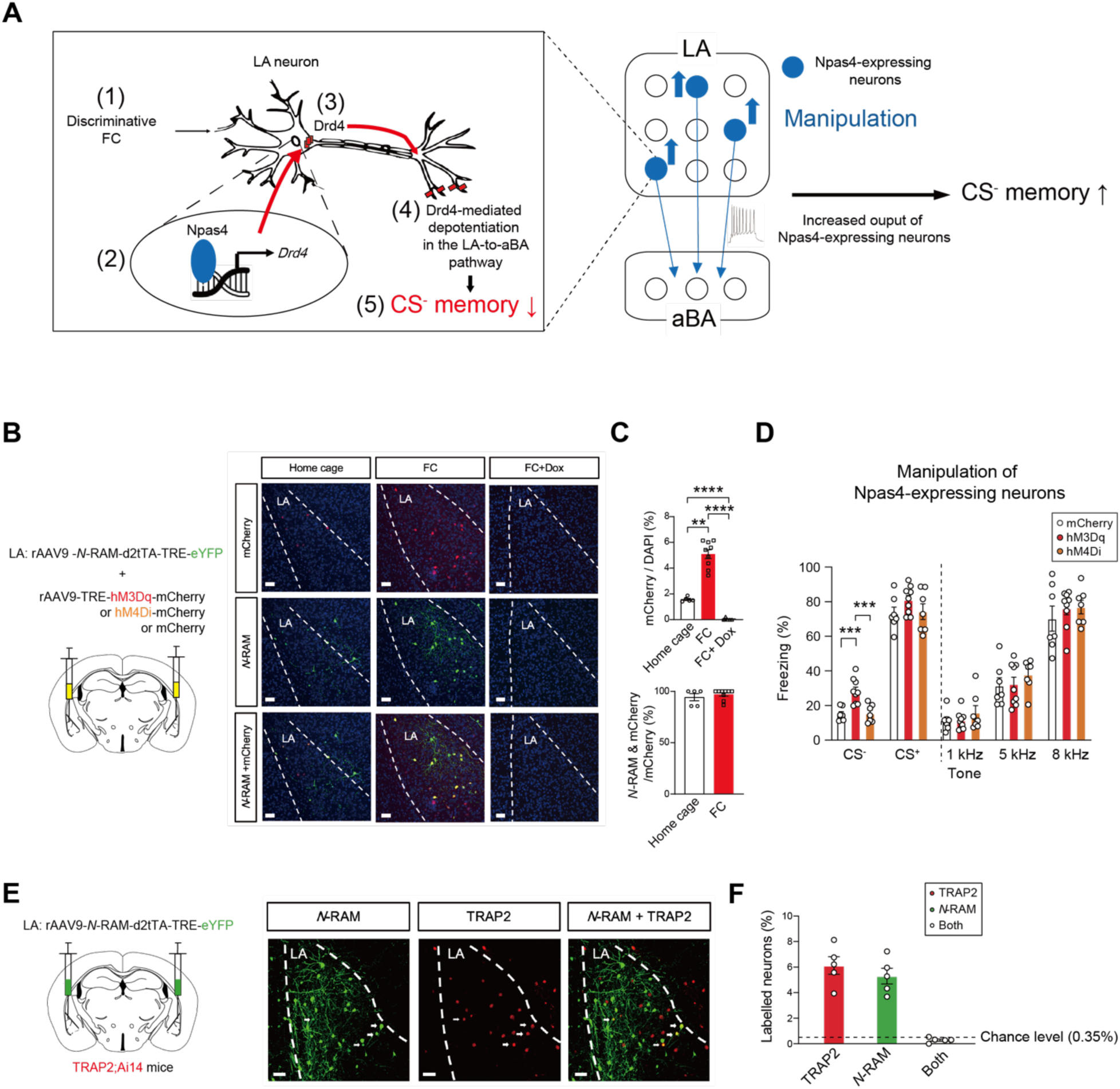
Retrievability of CS^-^ memory regulated by activity of Npas4-expressing neurons. **(A)** A descriptive diagram for Npas4-Drd4 signaling and manipulation of Npas4-expressing neurons which determine the expression of CS^-^ memory. **(B)** Schematic for labeling and activity manipulation of *N*-RAM-labelled neurons in the LA (left). Representative images for *N*-RAM labeling and DREADD expression within the LA areas outlined by dotted lines (right). Scale bars, 50 μm. **(C)** DREADD-expressing neurons relative to DAPI staining at each group (top, Home cage, n = 5; FC, n = 9; FC + Dox on, n = 6 mice; One-way ANOVA with Tukey’s *posthoc* test, \*\**P* = 0.0062, \*\*\*\**P* < 0.0001). Co-localization ratios of DREADD-expressing neurons denoted with mCherry and *N*-RAM-labelled neurons denoted with eYFP between home cage and FC groups (bottom, Home cage, n = 5; FC, n = 9 mice). **(D)** Activity manipulation of Npas4-expressing neurons in the generalization test (mCherry, n = 7; hM3Dq, n = 9; hM4Di, n =7 mice; One-way ANOVA with Tukey’s *posthoc* test, \*\*\**P* < 0.001). **(E)** Schematic for labeling of c-Fos- and Npas4-expressing cells in the LA (left). Representative images for engram neurons expressing either Npas4 or c-Fos after FC (right). White arrows indicate co-labelled neurons for c-Fos and Npas4. Scale bars, 50 μm. **(F)** Ratios of singularly- and co-labeled neurons within the LA (n = 5 mice). Data are shown as mean ± SEM,

Even a single memory episode from contextual fear conditioning can produce multiple neural engrams in the dentate gyrus, each of which has functionally and molecularly different features and functions (Sun et al., 2020). To this end, we sought to label these engram neurons activated by discriminative FC via the employment of two different approaches, *N*-RAM reporter and TRAP2 (targeted recombination in active population) mice (DeNardo et al., 2019). AAV encoding *N*-RAM probe was micro-infused into the LA of TRAP2-reporter mice (TRAP2; Ai14) (Figures 6E and Figure 6-figure supplement 1F): tdTomato expression was driven by the c-Fos promoter in the presence of 4-hydroxytamoxifen (4-OHT) while eYFP expression was driven by the *N*-RAM promoter in Dox-off period. This labeling experiment revealed two independent neural engrams with minimum overlapping (Figure 6F). If these two types of marked LA neurons exert separate functions, especially in the regulation of fear expression as previously shown in the dentate gyrus, Npas4-expressing engram neurons would intimately control the retrievability of CS^-^ memory through controlling the conversion of functional states of neural engrams for CS^-^ memory while c-Fos-expressing neurons could primarily underlie CS^+^ memory (Abdou et al., 2018). The balanced orchestration of two distinct neural engrams would maintain appropriate fear expression to ambient stimuli and help prevent the occurrence of pathological conditions such as PTSD.

## Discussion

The present study revealed distinct processing of irrelevant information for CS^-^ memory and the latent storage in amygdala circuits, which differed from those for CS^+^ memory. Retrievability of CS^-^ memory was gated by synaptic plasticity of specific amygdala circuits, the LA-to-aBA pathway. Stress exposure or CORT administration prolonged the retrievability of CS^-^ memory and modified neural representation of CS^-^ in aBA principal neurons. Npas4-mediated Drd4 synthesis during memory consolidation would help shift functional states of neural engrams for CS^-^ memory: arbitrary activation of the Npas4-expressing neuronal population in the LA is sufficient to convert “silent” CS^-^ memory engrams to “active” states. Although we focused on discriminative fear memory primarily regulated by the amygdala circuitry, the engram-mediated storage and state conversion of individual engrams may generally emerge in other brain areas and potentially could modulate various types of memories as well.

### Processing of information for CS^-^ memory in the amygdala circuits

Dissociating the stimuli predicting threats from others is an essential capability such that animals survive environmental circumstances for behavioral adaptation. Failure of such dissociation often heightens the sensitivity to irrelevant stimuli, which may cause fear-related diseases such as PTSD (van der Kolk, 1997; Pitman et al., 2012). Selecting appropriate information in ambient circumstances does not necessarily involve the removal of irrelevant information for CS^-^ memory, but it would rather serve as memory resources for future usage.

Sensory information for CS^-^ memory has been traditionally viewed as a safety signal. In discriminative FC, however, defensive responses to CS^-^ could arise in a “safety-by-comparison” way, different from those to safety cues in the learning: fear responses to CS^-^ become diminished by its comparison with CS^+^ (Genud-Gabai et al, 2013). CS^-^ memory could be retrieved at the STM time point but the freezing levels were significantly lower than those of CS^+^ (Figure 1), indicating that subject animal perceived CS^-^ as a relatively safer stimulus than CS^+^. Freezing levels to CS^-^ at the LTM point was further reduced after memory consolidation, consistent with the effectiveness of prior comparison between CS^-^ and CS^+^ onto fear expression across behavioral tests. Akin to the distinct processing for safety and CS^+^ memories (Rogan et al, 2005; Sangha et al, 2013), CS^-^ memory would be processed and stored, differently from CS^+^ memory. Accordingly, exposure to CS^+^ and CS^-^ drove divergent activity from LA and BA neurons after discriminative FC (Collins and Pare, 2000). In addition, the theta frequency range synchrony between the mPFC and the BLA was increased in the animals that were able to successfully distinguish CS^-^ from CS^+^. Thus, CS^-^ information could be represented and compared with CS^+^ likely in the amygdala circuits (Likhtik et al., 2014). Given our findings that specific memories are mediated by distinct engrams, it is possible that memory representation contained in each group of neural engrams could be assessed and compared through neuronal and circuit computation. The underlying circuit and molecular mechanisms of these interactions merit further investigation and should provide a refined understandings for the processing of CS^-^ memory and modulation of fear memory in general.

### Npas4-expressing neurons for processing and expression of CS^-^ memory

Neural engrams keeping memory should be reactivated for successful retrieval (Tonegawa et al., 2015). Forgetting, a reduction in retrievability of successfully encoded information (Tulving, 1974), would arise through the failure of engram reactivation as if silent neural engrams remained refractory upon exposure to stimuli (Ryan and Frankland, 2022). If certain memory was stored in silent neural engrams, those could be retrieved by artificial manipulation (direct optogenetic stimulation of cell bodies) (Josselyn and Tonegawa, 2020). Given the requirement of Npas4 and its downstream effectors such as Drd4 for the latent storage (could be artificially retrieved) of CS^-^ memory, *de novo* protein synthesis does not allow only for LTM of CS^+^ but also for processing and storage of CS^-^. As synaptic plasticity could occur exclusively in engram neurons (Zhou et al., 2009; Kim and Cho, 2017), the decremental fear responses to CS^-^ from the STM to LTM time points could reflect the state conversion of neural engrams for CS^-^ memory into the silent state through protein synthesis-dependent synaptic plasticity. The functionality conversion of neural engrams seemed to be an efficient means for appropriate memory operation.

It is intriguing that the expression of CS^-^ memory was mainly determined by the activity of the neuronal population marked by Npas4 whereas c-Fos-expressing neurons in the amygdala primarily mediated CS^+^ memory (Hall et al., 2001; Abdou et al., 2018). Npas4 is selectively induced by neuronal activity (Lin et al., 2008) and is involved in linking neural activity to memory discrimination (Sun and Lin, 2016), reminiscent with the regulatory roles of Npas4 for reduced expression of CS^-^ memory and conversion of the engram states. In fact, as Npas4 forms unique heterodimers (Brigidi et al., 2019; Sharma et al., 2019), it might trigger distinct activity-dependent gene programs which differed from the c-Fos-induced gene cascade (Yap et al., 2021). The distinct gene programs triggered by Npas4 and c-Fos would engender specialized behaviors depending on the involved IEG-dependent pathway. Interestingly, Npas4 regulated both excitatory and inhibitory synaptic transmission and plasticity in an activity-dependent manner (Bloodgood et al, 2013; Weng et al., 2018). The enhanced fear expression to CS^-^ that we observed with chemogenetic stimulation of Npas4-expressing LA neurons, supported the possibility that activation of the silent engram neurons enabled subject animals to re-access information for CS^-^ memory by affecting synaptic transmission and plasticity and thereby resulted in an elevation of the fear responses to CS^-^.

Although Npas4-expressing engram neurons are most likely to control and mediate the expression of CS^-^ memory, their cellular features that impart behavioral outcomes remained unknown. It was previously shown that engram neurons in the hippocampal CA1 region had lower spine density in retrograde amnesic than in non-amnesic ones (Roy et al., 2017). Accordingly, the reversal of spine density by expression of α-p-21-activated kinase 1 (PAK1) caused an increase in memory recall. For the long-term systems consolidation of fear memory, engram neurons in the prefrontal cortex should become functionally active concurrently with their increased spine density while the engram neurons in the hippocampus were inactive with the decreased spine density (Kitamura et al., 2017). Subsequent morphological examinations could provide systematic insights into cellular features of the engram neurons and potentially their state conversion for discriminative memory.

### Npas4-Drd4 axis controlled by CORT

The increasement of CORT levels results in maladaptive fear memory including impairment in stimulus discrimination (Klausing et al., 2020) and hypermnesia (Kaouane et al., 2012). We found that CORT could keep the synaptic transmission elevated and simultaneously prolong the retrievability of CS^-^ memory by affecting Npas4-Drd4 axis. It was widely documented that under stress exposure, CORT promoted synthesis of the target genes and memory retention for stressful episodes through binding to GR (de Quervain et al., 2016). The regulator regions of GR target genes, called glucocorticoid response elements (GREs), could be divided into several types: simple, half-site, negative, tethering and composite GREs (Schoneveld et al, 2004). A tandem repeat of half-site GREs would be negative GREs that inhibit target gene expression (Ou et al, 2001). Five putative tandem repeats of GRE half-sites are present in the *Npas4* promoter and deletion of three of them alleviated suppression of the promoter activity in the presence of CORT (Furukawa-Hibi et al, 2012). Therefore, Npas4-Drd4 axis could be affected by CORT through GR-mediated reduction of Npas4 production in a similar way.

The expression of Drd4 was reduced by CORT treatment in a neuronal cell line (Barros et al, 2003). However, how Drd4 expression was impeded by stress exposure and the resultant elevation of CORT levels remained unknown thus far. We took advantage of CUT&Tag assay and here provided the first experimental evidence for the interaction of Npas4 to the *Drd4* promoter in amygdala tissues. Combined with our luciferase results, CORT would reduce Drd4 expression through GR-mediated down-regulation of Npas4. The enhanced activity of Drd4 would result in depotentiation of the LA-to-aBA pathway of CORT-injected animals, which normalized the synaptic transmission and ameliorated the expression of CS^-^ memory. Synaptic depotentiation, or reversal of LTP (long-term potentiation), has been proposed to increase the capacity and efficiency for memory storage and processing (Huang and Hsu, 2001). Reversal of LTP in the hippocampal CA1 was observed after exposure to novel contexts (Xu et al, 1998) that promoted long-term memory (Moncada and Viola, 2007). Since activation of Drd4 caused the reversal of LTP (Kwon et al, 2008; Izumi and Zorumski, 2017), Npas4-mediated induction of Drd4 production could reverse LTP in the LA-to-aBA pathway and normalize the fear responses to CS^-^. Therefore, Npas4-Drd4 upregulation during memory consolidation is most likely to be a prerequisite for appropriate memory adaptation that was precluded by GR signaling.

This study reveals the unknown cellular and molecular mechanisms whereby irrelevant experience is processed and can induce abnormal fear responses after a stressful experience. We also provide the first experimental evidence that Npas4-expressing neurons could constitute silent engrams for CS^-^ memory and Npas4 regulates the synaptic transmission of the LA-to-aBA pathway in a Drd4-dependent manner, which demarcates the range of fear expression. Npas4 could be used as a valuable empirical means to capture CS^-^ memory-bearing engram neurons, which has never been attained thus far. Since excessive and aberrant expression of fear responses to ambient irrelevant stimuli is one of the core symptoms of fear-related disorders including PTSD, further elucidation of the molecular and circuit mechanisms underlying the processing of CS^-^ memory will open new avenues for the development of therapeutic treatments for PTSD.

## Materials and Methods

### Animals

All procedures performed for animal experiments were approved by the institutional animal care and use committee of Pohang University of Science and Technology (POSTECH), South Korea in accordance with the relevant guidelines. Mice were housed under a 12-hour light/dark cycle with *ad libitum* for water and food. Male adults of C57Bl/6J (Jackson Laboratory), C57Bl/6J-Tg(Rspo2-cre)Blto (RBRC10754, RIKEN, Japan), B6.Cg-Tg(Camk2a-cre)T29-1Stl/J (JAX #0053579), Fos^tm2.1(icre/ERT2)Luo^/J (JAX #030323), Ai14 (JAX #007914), *Drd4* knockout (JAX #008084) and Npas4^flx/flx^ mice (Lin et al., 2008) were used.

#### Construct subcloning

For the Npas4 expression vector, RNA extract from P5 C57Bl/6J brain was used to generate cDNA template for *Npas4*. The *Npas4* sequence (NM 153553.5) was amplified with PCR and inserted into pcDNA3.1 vector using KpnI and XbaI enzyme sites. The wild type (WT) *Drd4* promoter (ENSMUSR00000726369) was constructed using gDNA template extracted from C57Bl/6J brain. The *Drd4* promoter was subcloned into pGL3 Basic (Promega, E1751) vector using XhoI and HindIII enzyme sites. Two types of mutant *Drd4* promoters defective in Npas4 binding were constructed. The first mutant *Drd4* promoter was generated by deletion of two Npas4 consensus sequences (TCGTG) within the WT *Drd4* promoter region. The second mutant *Drd4* promoter was generated by deletion of the CUT&Tag-yielded interaction sequence containing two Npas4 motifs. *Drd4* promoters were subcloned and then used for the luciferase assay (Phusion^TM^ High-fidelity DNA polymerase, F530, Thermoscientific^TM^).

#### Virus procurement

*N*-RAM probe (pAAV-*N*-RAM-d2tTA-TRE-mKate2) was kindly provided by Dr. Yingxi Lin (Sun et al., 2020). Within the construct, mKate2 sequence was substituted with eYFP using NheI and AscI enzyme sites. Virus production was performed according to our established protocol (Kwon et al., 2015). In brief, pAAV-*N*-RAM-d2tTA-TRE-eYFP was co-transfected with AAV helper plasmids to HEK293T cells using Lipofector Q (AptaBio, South Korea). AAV particles were isolated and purified with iodixanol-gradient ultracentrifugation. The lysate was washed and concentrated with Amicon Filter (Millipore) to achieve at least 2.0 x 10^12^ gc/ml. The other viruses were obtained from Addgene, University of North Carolina (UNC) Vector Core and BrianVTA (China) unless described otherwise: rAAV9-CaMKII*α-*hChR2(*H134R*)-eYFP (26969) and rAAV5-Syn-Flex-GCaMP6f-WPRE (100833) from Addgene; rAAV5-CaMKII*α-*mCherry-Cre, rAAV5-CaMKII*α*-mCherry and rAAV9-CaMKIIα-eYFP from UNC vector core; rAAV9-TRE3g-hM4Di-mCherry, rAAV9-TRE3g-hM3Dq-mCherry and rAAV9-TRE3g-mCherry from BrainVTA; rAAV5-cKD-scRNA (scrambled RNA, 5’-GCACTACCAGAGCTAACTCAGATAGTACT-3’)-eYFP and rAAV5-cKD-shDrd4-eYFP (Kwon et al., 2015).

#### Stereotaxic surgery

For micro-infusion of viruses, mice were anesthetized by intraperitoneal (I.P.) injection of ketamine and xylazine mixture (100 mg/kg and 14 mg/kg, respectively). Mouse head was fixed with stereotaxic apparatus (Kopf Instrument). Coordinates for virus injection were as follows: -1.6 mm anteroposterior (AP), ±3.4 mm mediolateral (ML), and -4.3 mm dorsoventral (DV) for the lateral amygdala (LA); -1.0 mm AP, ±3.2 mm ML, and -5.0 mm DV for the anterior basolateral amygdala (aBA). The total injection volume was 4.6 – 9.2 nl for injection for the LA and approximately 300 nl for the aBA. AAV-containing solution was infused at 23.0 nl/sec using a Nanoject II or III (Drummond). Fiber optic cannulas (200 μm diameter, 0.22 NA, Newdoon Technologies, China) were bilaterally implanted just above the aBA. Guide cannula (C315G/SPC, P1 Technologies) was also bilaterally implanted above the LA.

#### *Ex vivo* electrophysiology

Acute brain slices were prepared in coronal sections. All the chemicals used for electrophysiological experiments were from Sigma unless otherwise specified. First, principal neurons in aBA were identified with their large soma size and capacitance exceeding 100 pF. Excitatory postsynaptic currents (EPSCs) were recorded from the aBA principal neurons in amygdala brain slices (-0.82 to -1.22 mm from bregma). tdTomato fluorescent signals were visualized with an upright microscope equipped with standard epifluorescence. Whole-cell patch clamp recordings were made with a MultiClamp 700B amplifier (Molecular Devices) in artificial cerebrospinal fluid (aCSF) containing 119 mM NaCl, 2.5 mM KCl, 2 mM MgSO4, 1.25 mM NaH2PO4, 26 mM NaHCO3, 10 mM D-glucose and 2.5 mM CaCl2, equilibrated with 95% O2 and 5% CO2 (pH 7.3 - 7.4) at room temperature (RT). Recording electrodes (8-10 MΩ) were filled with internal solution containing 130 mM CsMeSO4, 8 mM NaCl, 10 mM phosphocreatine,10 mM HEPES, EGTA, 2 mM, 0.5 mM MgATP, 0.1 mM NaGTP, and 5 mM QX-314, at pH 7.2, adjusted with CsOH (Kwon et al., 2015) in a voltage-clamp configuration. Recording of EPSCs was made at -70 mV holding potential in presence of 100 μM picrotoxin while series resistance (10 - 30 MΩ) was monitored. Monosynaptic natures of excitatory transmission evoked in the LA- to-aBA pathway was tested by bath application of tetrodotoxin (TTX, 1 μM, Tocris) and subsequently 4-AP (400 μM). To ascertain the excitatory synaptic transmission of the LA-to-aBA pathway, NBQX (10 μM, Tocris) and APV (50 μM) was applied in the absence of PTX. We also perfused a Drd4 agonist (PD-168,077, 200 nM, 1065, Tocris), Drd1/Drd5 agonist (SKF 38393, 10 μM), or Drd4 antagonist (L-745,870, 50 nM, Tocris) to examine the dopaminergic signaling. To assess the pre/postsynaptic contribution of Drd4 for alteration of synaptic transmission, guanosine 5’-[β-thio]diphosphate trilithium salt (GDPβS, 0.5 mM) was included in the internal solution.

For ratio measurement of AMPAR/NMDAR-EPSCs (α-amino-3-hydroxy-5-methyl-4-isoxazolepropionic acid receptor/N-methyl-D-aspartate receptor-mediated EPSCs), we first elicited AMPAR-EPSCs at -70 mV (10 traces) and then NMDAR-EPSCs at +40 mV holding (10 traces) with the delivery of 473 nm light pulses (0.5 ms, 0.05 Hz) in presence of picrotoxin (PTX, 100 μM). The amplitudes of NMDAR-EPSCs were obtained 50 ms after stimulus onset while those of AMPAR-EPSCs were detected at their peaks. To construct the input-output curves of EPSCs, light-evoked EPSC was measured at -70 mV, five of which were averaged at each light power. For measurement of paired-pulse ratios (PPRs), ten ratios of 2^nd^ EPSC / 1^st^ EPSC were averaged at interstimulus intervals (ISI) starting from 50, 100, 150 to 250 ms at -70 mV holding. For progressive blockade of NMDAR-EPSCs (Kim and Cho, 2020), NMDAR-EPSCs were recorded at +40 mV with the presence of NBQX (10 μM) and PTX (100 μM). The amplitudes of NMDAR-EPSCs were adjusted to 150 – 200 pA with adjustment of light power. After obtaining the stable baselines, we bath perfused MK-801 (10 μM, Tocris) for 15 minutes while 60 NMDAR-EPSCs were elicited every 10 sec. The decay time constant (τ) was calculated with the formula, *I*(*n*) = *I*1 *exp*(−*n*/τ) (n, stimulus number; *I*(*n*), the peak amplitude of *n*th NMDAR-EPSC; *I*1, the peak amplitude of the first NMDAR-EPSC).

To validate efficacy of the used DREADDs, whole-cell recordings were made using recording electrodes (8-10 MΩ) filled with the internal solution containing 120 mM potassium-gluconate, 5 mM NaCl, 0.2 mM EGTA, 1 mM MgCl2, 10 mM HEPES, 2 mM MgATP, and 0.2 mM NaGTP. Then, membrane excitability was measured in current-clamp configuration by applying step currents (Δ50 pA, 1000 ms) from –200 pA to +200 pA. Rheobase was determined as the lowest injected current that started to evoke an action potential in 2 pA resolution. Resting membrane potentials (RMPs) were also measured after stabilization of baselines (typically 10 minutes). RMPs and rheobases were again obtained after perfusion of a DREADD agonist compound 21 (C21, 10 nM, HB6124, Hello Bio Ltd). Measured RMPs were averaged for 5 minutes before and after C21 application.

Two protocols were established in either the presence or absence of PTX, respectively for optical induction of long-term depression (LTD). The sub-threshold LTD protocol consisted of 473 nm light at 0.5 Hz (150 pulses of single pulses of 0.5 ms duration). The *in vivo* LTD protocol was established in the absence of PTX to recapitulate *in vivo* circumstances. The *in vivo* LTD protocol consisted of the delivery of 450 pulses of 473 nm light at 0.5 Hz. Those protocols were *ex vivo* tested for LTD induction only once per each brain slice.

#### *In vivo* LTD induction and drug infusion

For *in vivo* induction of LTD in the LA-to-aBA pathway of behaving animals, the LTD protocol (450 pulses at 0.5 Hz; duration of each pulse; 473 nm at 2 - 4 mW) that had been approved through *ex vivo* and *in vivo* recording was delivered bilaterally through optic fibers implanted above the aBA following micro-infusion of either rAAV9-CaMKIIa-hChR2*(H134R)*-eYFP or rAAV9-CaMKIIa-eYFP into the LA. Optic cables kept disconnected for 15 minutes after LTD induction, and 45 minutes later, freezing responses were monitored with the presentation of paired conditioned stimuli (CS^+^) or unpaired conditioned stimuli (CS^-^).

For Drd4 agonist infusion into the LA, mice received either 300 μl of PD-168,077 (200 nM, Tocris) or vehicle (Dimethyl sulfide, D2650, Sigma) through internal cannula (C315I/SPC, P1 Technologies) using Pump 11 Elite (Harvard Apparatus) at a rate of 0.1 μl/minute. Mice were kept with internal cannula for additional 10 minutes and remained at home cage for 30 minutes before the cue test. The subject mice were cardially perfused immediately after behavioral tests for histological verification of cannula implantation sites.

#### Behavioral tests

For fear conditioning, three different chambers (26 cm x 26cm x 24 cm, length x width x height) were used such that mice could distinguish contexts: context A was made of black opaque PVC walls and a grid floor, which was cleaned with 70% ethanol before each trial; context B was made of transparent plastic walls and a white PVC floor covered with sawdust bedding and was scented with peppermint odor; context C was made of black PVC walls with PVC floor covered with corncob bedding. All the chambers were situated in a sound attenuation box in which was equipped with an infrared beam as well as a USB camera (Sentech, Japan) connected to a personal computer. Animal behaviors, including freezing and locomotion, were acquired and analyzed with ExthoVision XT (Noldus, Netherland)

Mice underwent 2 minutes of acclimation in the context A. A tone (CS^+^: 10 kHz, 75 dB, 30 sec) was co-terminated with electric foot shocks (unconditioned stimulus, US: 0.4 mA, 0.5 sec). A tone (CS^-^: 2 kHz, 75 dB, 30 sec) was delivered without application of electric foot shocks. For discriminative fear conditioning, 8 pairings of CS^+^-US and 8 CS^-^ were presented in pseudorandom intertrial intervals (varied from 45 to 90 sec). Either CORT (Corticosterone HBC complex, 5 mg/kg, C174, Sigma) or saline (0.9% NaCl in distilled water) was I.P. injected to mice immediately after completion of fear conditioning as previously described (Kaouane et al., 2012). For CS^+^_only_ group, all procedures were identical except the absence of CS^-^ presentation. To ascertain the efficacy of a glucocorticoid receptor (GR) antagonist, either mifepristone (10 mg/kg, M8046, Sigma) or vehicle (Dimethyl sulfide, 0.16 ml/kg, D2650, Sigma) was injected to mice immediately after fear conditioning

For the cue test, mice were initially placed in context B for 2 minutes of acclimation 24 hours after fear conditioning, and then CS^-^ was presented 3 times in pseudorandom intertrial intervals (varied from 45 to 90 sec) followed by 3 CS^+^ in the same pseudorandom intervals. For the generalization test, subject mice underwent the same procedure as in the cue test, but now tones of various frequencies were presented as well. The tones starting from 1, 2, 5, 8, to 10 kHz (75 dB for 30 sec each) were applied (3 times at each frequency in the same range of pseudorandom intervals). To switch tone frequencies for CS^+^ and CS^-^ (to test the sound-tuning effect), all the procedures were identical, but 2 kHz and 10 kHz were used for CS^+^ and CS^-^, respectively. In the generalization test using switched frequencies, tones starting from 12, 10, 8, 5 to 2 kHz were delivered (3 times at each frequency in the same range of pseudorandom intervals). Freezing responses were assessed by measuring the time duration for absence of movements except respiratory activity. The discrimination index was calculated as [(CS^+^ freezing – CS^-^ freezing) / (CS^+^ freezing + CS^-^ freezing)] (Kim and Cho., 2017).

For identification of brain regions involved in the processing of memory information for CS^-^ or CS^+^ cues, animals underwent the identical cue test 24 hours after fear conditioning, but either CS^-^ (2 kHz, 75 dB, 30 sec) or CS^+^ (10 kHz, 75 dB, 30 sec) was presented 3 times. 90 minutes after behavioral tests, subject mice were sacrificed for histological analysis.

Subject animals underwent the elevated plus maze (EPM) test with the maze consisting of two open arms (50 cm x 10 cm, length x width) and two closed arms (50 cm x 10 cm x 30 cm, length x width x height, black wall) that were elevated at 60 cm above the ground. On the subsequent day, the open field test (OFT) was performed in a rectangular box (60 cm x 40 cm x 30 cm, length x width x height). Mice were placed for 15 minutes to assess free exploration of EPM and OFT. For the pain threshold test, mice were exposed to electric shocks with a sequence of 0.05, 0.08, 0.1, 0.15, 0.2, and 0.25 mA for 0.5 sec each (with 60 sec interval) after 2 minutes of acclimation in a conditioning chamber. The pain threshold was determined at the intensity of the electric shock at which subject mice started to jump or scream. For exposure to restraint stress, the mice were immobilized in a ventilated Plexiglas tube for 2 hours and then returned to home cage.

#### Measurement of serum CORT

Immediately after fear conditioning, subject animals were rapidly decapitated, and their blood was collected. The blood was incubated for 20 minutes at RT and centrifuged in 2000 r.c.f. for 10 minutes at 4°C to collect serum (stored at – 80°C). All samples underwent only one freeze-thaw cycle. CORT ELISA kits (ADI-900-097, Enzo Life Sciences) were used following the manufacturer’s instructions. The samples were thawed immediately prior to actual measurement. The dilution factor was 40-fold. The plates were red on a 96 Plate Reader (Enzo Life Sciences).

#### Labeling and manipulation of engram neurons

To label Npas4-expressing neurons and for activity manipulation, wild type mice were fed with doxycycline (Dox)-containing diet (60 mg/kg) before virus injection. Virus cocktail of rAAV9-*N*-RAM-d2tTA-eYFP and one out of rAAV9-TRE3g-hM3Dq-mCherry or rAAV9-TRE3g-hM4Di-mCherry or rAAV9-TRE3g-mCherry (ratio of 1:3) was injected to the LA. 7 days after virus infusion, subject mice underwent 2 day-long Dox-off period. After 1 day-long Dox-off period preceded by fear conditioning, C21 (0.5 mg/kg) was I.P. injected 1 hour before behavioral tests.

For double-labeling of c-Fos- and Npas4-expressing neurons, TRAP2 Fos^tm2.1(icre/ERT2)Luo^/J; Ai14 double transgenic mice were generated and received virus injection of rAAV9-*N*-RAM- d2tTA-eYFP with an identical schedule as described above except of 4-OHT injection (50 mg/kg, 4-Hydroxytamoxifen, H7904, Sigma) immediately after fear conditioning. Subject animals were sacrificed for histological examination 7 days after fear conditioning. The chance level of co-expression of c-Fos- and Npas4-expressing neurons was determined with (eYFP / NeuN) x (tdTomato / NeuN) for each brain slice.

#### Quantitative Real-Time PCR

0.5, 4.5, 12 and 24 hours after fear conditioning, brains were collected and placed in ice-chilled Adult Mouse Brain Slicer Matrix (BSMAS005-1, ZIVIC Instruments). The coronal sections were made in 1 mm thickness. Amygdala regions were isolated with a Biopsy punch (15110-15, Ted Pella Inc). Total RNA was extracted using RNA extraction kit (Intron). 800 ng of extracted RNA was used for reverse transcription with iScript Reverse Transcriptase (BioRAD). The fold change of various genes was calculated through the delta-delta CT method using glyceraldehyde 3-phosphate dehydrogenase (GADPH) as a reference gene (Livak and Schmittgen., 2001). RNAs isolated from mice of home cage group were used as references for multiple comparisons.

Used primers as follows, Npas4: forward 5’-CTGCATCTACACTCGCAAGG -3’, reverse 5’-GCCACAATGTCTTCAAGCTCT-3’; Arc: forward 5’-TACCGTTAGCCCCTATGCCATC-3’, reverse 5’-TGATATTGCTGAGCCTCAACTG-3’; c-Fos: forward 5’-ATGGGCTCTCCTGTCAACACAC-3’, reverse 5’-ATGGCTGTCACCGTGGGGATAAAG-3’; GAPDH: forward 5’-CA TGGCCTTCCGTGTTCCT-3’, reverse 5’-TGATGTCATCATACTTGGCAGGTT-3’ (Ramamoorthi et al., 2011); Drd4: forward 5’-GCCTGGAGAACCGAGACTATG-3’, reverse 5’-CGGCTGTGAAGTTTGGTGTG -3’ (primerBank ID 6681223a1).

### *In vivo* Ca^2+^ imaging from individual neurons

One week after virus infusion (rAAV5-Syn-Flex-GCaMP6f-WPRE), a Proview lens (0.6 mm diameter, Inscopix) was implanted above the aBA. Before lens implantation, custom-made needles (21 G) stayed 15 minutes to keep piercing the brain tissues. The lenses were lowered at a rate of 0.03 mm/sec. The top side of lenses was sealed with Silicone Elastomer (Kwik-CastT^M^, WPI). 2 weeks after lens implantation, the baseplate was installed to hold the micro-endoscope

*In vivo* Ca^2+^ imaging was performed typically 1 - 2 weeks after baseplate installation. Mice were briefly anesthetized with isoflurane to assemble/disassemble the micro-endoscope (nVoke2, Inscopix). The mice stayed in the home cage for at least 1 hour after assembling of micro-endoscope for the recovery. Imaging parameters including frame rates (10 Hz), gain (1 -2.5), LED powers and focal planes were adjusted and set constant for each subject throughout the entire imaging sessions.

For data processing and analysis, we used IDPS software (Inscopix) for motion correction and exported them into a single image stack file in a TIFF format. Constrained non-negative matrix factorization (CNMF-E) (Zhou et al., 2018) was utilized for extraction of fluorescence dynamics at the single cell level and spatially down-sampled with factor 4.

To determine whether given responses from individual neurons were significantly modulated by tone presentation, we first collected a scaled version of ΔF/F (neuron.C_raw) around tone onset (10 sec before and after of tone onset). ΔF/F was down-sampled to a time bin of 1 sec. To overcome possible trial-to-trial variability, ΔF/F obtained from each trial was concatenated into before and after matrix, respectively. The neurons were defined as “responsive” if two matrices significantly differed (Wilcoxon signed-rank test, p <0.05) and as “non-responsive (NR)” if two matrices were found similar.

Out of those responsive neurons, we classified a group of neurons as “Excited” if the mean ΔF/F of after-matrix were higher than that of before-matrix. A group of neurons was classified as “Inhibited” if the case was the opposite. For further analysis, denoised ΔF/F was converted to z-scored activity. The z-scores activity of each neuron was obtained with (F_t_ – F_m_) / SD (where F_t_ is the ΔF/F at time point t, F_m_ and SD are the mean and standard deviation of entire session, respectively). To compare the spatial distribution of given-classified neurons, we calculated the pairwise distance of each neuron’s centroid position within each group. Averaged cue response from each neuron was obtained as the mean of z-scored activity for 10 sec from the stimulus onset.

For population vector analysis, we down-sampled temporal z-scored activity to a time bin of 1 sec. From each mouse, we calculated n-dimensional activity vectors from individual neurons at each time bin in which the total number of extracted neurons was n (Grewe et al., 2017). We used principal component analysis (PCA) for dimensionality reduction of n-dimensional activity vector and projected them to 3-dimensional space composed of PC1, 2 and 3. Mean neural trajectory was obtained by averaging values from trials in response to each tone. The Mahalanobis distance (MD) was calculated between two different stimuli for comparison: MD^2^ = *(A-B)^T^ ⋅ Σ^-1^(A-B)* where *A* and *B* were individual averaged neural trajectory evoked by tones of different frequencies, and *A^T^* and *B^T^* were their transposes. The deviation of population vector distance (ΔPVD) was calculated by subjecting MD between two different tones from stimulus onset to all the time points. Mean ΔPVD was calculated for 30 sec from stimulus onset. To gain insights into correlation with behavioral data, averaged freezing magnitudes to CS^-^ were plotted with the mean ΔPVD between CS^-^ and CS^+^.

### *In vivo* electrophysiology

Custom-made optrodes were used for *in vivo* recording of local field potentials (LFPs). Electrode bundles consisted of 16 individually insulated nichrome wires (15-mm diameter, impedance of 80 - 300 KU, A-M systems) were fixed to a fiber optic cannula (200 μm diameter, 0.22 NA, Newdoon Technologies, China). Optrode bundles were attached to one 36-pin male nano dual row connector (Omnetics).

One week after virus infusion into the LA (rAAV9-CaMKII*α*-hChR2(H134R)-eYFP), optrodes were implanted just above the aBA using stereotaxic apparatus (Kopf Instrument). Connectors were referenced and grounded via 4 silver wires (127 μm diameter, A-M Systems) placed above the cerebellum. Implanted optrodes and nano-connectors were fixed with super-bond cement (Sun Medical).

While briefly anesthetized with isoflurane, electrodes were connected to a Cereplex μ headstage (Blackrock microsystems). Subject mice stayed in the home cage for at least 1 hour for recovery. While the headstage was connected to CerePlex Direct, local field excitatory postsynaptic potentials (fEPSPs) were acquired by 30 kilo-samples per second and bandpass-filtered between 10 and 1000 Hz in the home cage. 473 nm DPSS laser (Shanghai Laser & Optics Century, China) was used for optic stimulation (0.033 Hz; duration of single pulse, 5 ms) through Master-8 stimulator (AMPI, Israel). Laser power was adjusted to 30 - 40 % of which yielded maximum fEPSP amplitudes. fEPSP slopes were calculated in (Maximum fEPSP value – Minimum fEPSP value) x 0.8 / (decay time from 80 to 20 % fEPSP values) and averaged them for 5 minutes. fEPSPs were acquired from each animal for 4 sessions: before discriminative FC (for 1 hour, Before FC) and ① for the first 1 hour in the next day; ② for another 1 hour after injection of PD-168,077 or vehicle; ③ for the subsequent 1 hour. For *in vivo* validation of the LTD protocol, all the procedures were identical but substituted Drd4 agonist injection to LFS (450 pulses at 0.5 Hz; duration of each pulse; 5 ms). The recording sites was electrically lesioned for verification of optrode placements.

### Immunohistochemistry (IHC)

Mice were transcardially perfused first with phosphate-buffer saline (PBS) and then with 4 % paraformaldehyde (PFA) in PBS. Isolated brains were kept in 4 % PFA-containing PBS overnight at 4 °C. For cryosection, brains were first dehydrated in 30 % sucrose-containing PBS for 2 days at 4 °C. Dehydrated brains were frozen in 2 methyl-butane chilled with dry ice and then stored at - 80°C. The brains were sliced into 40 μm coronal sections using a Cryostat (Leica, Germany). The brain slices underwent blocking either with 4 % normal donkey serum (NDS) or normal goat serum (NGS) in 0.3 % Triton X-100-containing Dulbecco’s phosphate-buffered saline (DPBS) for 1 hour at RT. Subsequently, those slices were incubated at 4 °C for 24 hours in 1% NDS or NGS in 0.3 % Triton X-100 DPBS containing goat anti-Drd4 (1:500, sc-31481, Santa Cruz), rabbit anti-c-Fos (1:500, sc-52, Santa Cruz), rabbit anti-Npas4 (1:200, Activity Signaling), or mouse anti-NeuN (1:500, Sigma-Aldrich). Those slices were washed 3 times in DPBS (10 minutes at RT). After washing, they were incubated for 24 hours at 4 °C with Alexa Fluor 594-conjugated donkey anti-goat IgG (1:300, A-11058, Invitrogen), Cy3-conjugated goat anti-rabbit IgG (1:500, 111-165-144, Jackson ImmunoResearch laboratories Inc) or Cy5-conjugated goat anti-rabbit IgG (1:200, 111-175-144, Jackson ImmunoResearch laboratories Inc) in 1% NDS or NGS in DPBS containing 0.3 % Triton X-100. They were again washed 3 times in DPBS (10 minutes at RT) and then mounted with UltraCruz mounting medium (Santa Cruz) for imaging.

### RNAscope^®^ fluorescence in situ hybridization (FISH)

RNAscope^®^ multiplex fluorescent assay (320850, Advanced Cell Diagnostics) was used to visualize *Drd4* mRNA (Probe 418171), following the manufacturer’s instructions. Briefly, fresh frozen brains were sectioned into 15 μm slices using a Cryostat (Leica). Those slices were mounted to slide glasses and underwent fixation, dehydration, hydrogen peroxide, protease treatment steps. Finally, the *Drd4* probe was incubated for 2 hours at 40°C.

### Confocal imaging and analysis

Confocal laser scanning microscopes (FV-3000, Olympus, Japan) was used to acquire fluorescence images with objective lenses (UPLSAPO 20 X, 40 X, 60 X, Olympus). MetaMorph 7.7 (Molecular Devices) was used for quantitative analysis of Drd4 puncta. Numbers of DAPI staining, NeuN-, c-Fos- and Npas4-expressing cells were manually counted.

### Immuno-electron microscopy

Immuno-electron microscopic analysis was made in accordance with the previously established protocols (Masugi-Tokita and Shiegemoto, 2007; Kwon et al., 2015). To label axon terminals originated from the LA, we injected rAAV9-CaMKIIa-hChR2*(H134R)-*eYFP into the LA and kept them for 3 weeks in the home cage. After cardiac perfusion, the brain was sliced into 200 µm. Those aBA-containing amygdala sections were dehydrated in 10% sucrose solution in PBS. High-pressure freezing instrument (HPM 100, Leica) was used to preserve membrane structure. Then, the tissues were plunged in acetone and embedded in resin (Lowicyrl HM20, Electron microscopy sciences) at - 45°C for 2 days and UV-polymerized for 1 day in EM AFS2 (Leica). The aBA-containing polymerized blocks were further sliced into 100 nm thickness by an ultra-microtome (Leica), and slices were put on the Nickel grids (FCF200-Ni, Electron microscopy sciences). For immunolabeling, rabbit anti-GFP (1:20, LF-PA0046, AbFrontier, South Korea) and goat anti-Drd4 (1:20, sc-31481, Santa Cruz) antibodies were used as primary antibodies. They were secondarily labeled with donkey anti-rabbit antibody conjugated with 18-nm gold particle (711-215-152, Jackson ImmunoResearch) or donkey anti-goat antibody conjugated with 12-nm gold particle (705-205-147, Jackson Immuno Research), after blocking with 0.2% NDS-containing detergent-free PBS at 4°C overnight. We conducted post-staining with uranyl acetate (for 4 minutes) and Reynolds (for 2 minutes) to enhance the contrast. Images were obtained with a transmission electron microscope (JEM-1011, Jeol, Japan). Asymmetric synapses were defined by the presence of thick postsynaptic density (PSD) greater than 40 nm.

### CUT&Tag experiments

CUT&Tag assays were carried out as previously described (Kaya-Okur et al., 2019) with minor modifications. Briefly, amygdala tissues of three mice were homogenized with Dounce homogenizer in NE buffer (20 mM HEPES-KOH, pH 7.9, 10 mM KCl, 0.5 mM Spermidine, 0.1% Triton X-100, 20% Glycerol and 1x GenDEPOT Xpert protease inhibitor cocktail solution) and pooled to eliminate potential variances. Nuclei from tissues were captured with CUTANA™ Concanavalin A Conjugated Paramagnetic Beads (EpiCypher) and incubated with the primary antibody to Npas4 overnight on a nutator. Subsequently, the secondary antibody was added and incubated for 30 minutes at RT on a nutator. Then, unbound antibody was washed away with Digitonin150 buffer (20 mM HEPES-NaOH pH 7.5, 150 mM NaCl, 0.5 mM Spermidine, 0.01% digitonin and 1x GenDEPOT Xpert protease inhibitor cocktail solution). 20x CUTANA™ pAG-Tn5 (EpiCypher) was added to pooled nuclei such that the final concentration of the Tn5 was 1x in Digitonin300 buffer (20 mM HEPES-NaOH pH 7.5, 300 mM NaCl, 0.5 mM Spermidine, 0.01% digitonin and 1x GenDEPOT Xpert protease inhibitor cocktail solution). Nuclei with Tn5 were incubated for 1 hour at RT for Tn5 to bind to the primary and secondary antibodies. The nuclei were washed again with Digitonin300 buffer and incubated in Tagmentation buffer (20 mM HEPES-NaOH pH 7.5, 300 mM NaCl, 0.5 mM Spermidine, 10mM MgCl2 and 1x GenDEPOT Xpert protease inhibitor cocktail solution) for 1 hour at 37°C to activate tagmentation. Tagmentation buffer was then replaced with SDS release buffer (10 mM TAPS pH:8.5, 0.1% SDS) for 1 hour at 58 °C to release the tagmented DNA fragments. Lastly, SDS Quenching buffer (0.67% Triton-X 100) was added to neutralize SDS that could potently inhibit PCR. The following antibodies were used: primary antibody, rabbit anti-Npas4 (Activity signaling) and secondary antibody, goat anti-rabbit antibody for CUTANA™ CUT&Tag Workflows (13-0047, EpiCypher).

### Library preparation and sequencing for CUT&Tag

CUT&Tag library construction was performed using CUTANA™ High Fidelity 2X PCR Master Mix (15-1018, EpiCypher) following the manufacturer’s instruction. Primers and the tagmented chromatin were added to the master mix to achieve high fidelity amplification of NGS libraries. Tagmented chromatin was amplified by 18 cycles of PCR. The amplified DNA library was size selected (230 to 470 bp), measured by Qubit fluorometer and proceeded to sequencing. CUT&Tag libraries were sequenced on Illumina HiSeqX instrument with 150 bp paired-end reads according to the manufacturer’s instructions (Illumina).

### Data processing for CUT&Tag dataset

Low quality and adapter sequences were trimmed using ‘Trim Galore’ with ‘-q 30 --paired’ parameters. Trimmed reads were aligned using ‘Bowtie2’ (Langmead and Salzberg, 2012) for mm10 reference FASTA with ‘--local --very-sensitive-local --no-mixed --no-discordant’ parameters. Aligned reads were sorted using ‘Sambamba’ (Tarasov et al., 2015). For peak calling, ‘MACS2’ (https://github.com/macs3-project/MACS) was used with ‘callpeak --keep-dup all --p 1e-5 --f BAMPE’ parameters. To filter blacklist regions, overlapped peaks with blacklist regions were removed using ‘bedtools intersect’ (Quinlan and Hall, 2010). Blacklist regions were downloaded from ENCODE (https://www.encodeproject.org/references/ENCSR094CNP/). Filtered peaks were annotated by nearest genes from the peak center using ‘annotatePeaks.pl’. The promoter region was defined within 2 kb from TSS.

Before making bigWig track files, BAM files were converted to BED format using ‘bedtools bamtobed’ and tag directories were made using ‘makeTagDirectory’ with default options. UCSC genome browser was used for genome wide visualization with bigWig files and called peaks. For making bigWig track files, ‘makeUCSCfile’ in HOMER package was used with ‘-- norm 1e7’ parameter. Heatmap analysis was performed using ‘Deeptools’ package (Ramirez et al., 2016).

The CUT&Tag data discussed in this publication have been deposited in NCBI’s Gene Expression Omnibus (GEO) and are accessible through GEO series accession number GSE194069.

### Luciferase analysis

Luciferase assay was performed using Dual Luciferase Assay System (E1910, Promega). Following the manufacturer’s guideline, HEK-293T cells were grown in 12-well plates and transfected with the constructs (either *Npas4*-subcloned pcDNA3.1 vector or *Npas4*-empty pcDNA 3.1 vector and pGL3 Basic vector containing either WT or mutant *Drd4* promoter) and polyethylenimine (PEI). Passive Lysis Buffer was added to each well (100 μl) for detachment of cells. The cell lysate was centrifuged for 30 sec at 12,225 r.c.f. and supernatant (40 μl) was transferred to 96-well white plates (136101, Nunc). Luciferase assay buffer 2 (50 μl) was added to each well. Firefly luciferase activity was first measured for 10 sec and then Stop & Glo Buffer (Promega) was added. Subsequently, Renilla luciferase activity was measured for 10 sec. The injection of buffer and luminescence detection was conducted by an automated injector and plate reader (Infinite M200 PRO, TECAN).

### Statistical analysis

Statistical tests were conducted using Prism 8 (GraphPad, USA). The statistical details were specified in supplementary table 1. The datasets were first tested for Gaussian distributions with a Shapiro-Wilk normality test. When normal distribution was not rejected in this way, the datasets were compared with Student’s t-test or ANOVA test but otherwise, were tested with Mann-Whitney U-test. To measure the association between two variables, Pearson’s correlation coefficient was calculated. All data are presented as mean ± standard error of the mean (SEM) in figures. The p-value threshold of significance was set at 0.05, presented as *p < 0.05, **p < 0.01 or *** p < 0.001. Statistical significance is indicated using symbols (* and #).

## Author contributions

B.K. and J-H.K. conceived the study and designed the experiment. B.K. and J.-Y.Y. performed *ex vivo* recording and behavioral experiments. B.K. performed *in vivo* Ca^2+^ imaging and W.C. wrote the code for analysis of the data. B.K. and R.D. performed CUT&Tag experiments and D.U. analyzed the data. K.S. performed and analyzed *in vivo* LFP recording experiment. S.B.L. performed luciferase assay. S.B.L., T.Y. and S.T.B. designed and subcloned DNA constructs. B.K. and T.Y. performed CORT level measurement. H.J.K. performed immune-EM experiments. B.K., R.D., T.Y., S.L., D.U., S.K.P., T.-K.K., S.-B.P. and J.-H.K. wrote the article. J.-H.K. supervised the entire work.

## Acknowledgments

We appreciate Crag H. Bailey (Columbia) and Eric R. Kandel (Columbia) for critical reading and helpful suggestions throughout the entire study. We also thank Michael E. Greenberg (Harvard) for providing Npas4^flx/flx^ mice and reagents, and Yingxi Lin (SUNY Upstate) for the N-RAM construct. This work was supported by the National Research Foundation of Korea (NRF) (2018R1A3B1052079 and 2018M3C7A1024152 to J.-H.K.).

## Declaration of Interests

The authors declare no competing interests.

## Supplemental information titles and legends

**Figure 1 – figure supplement 1.**
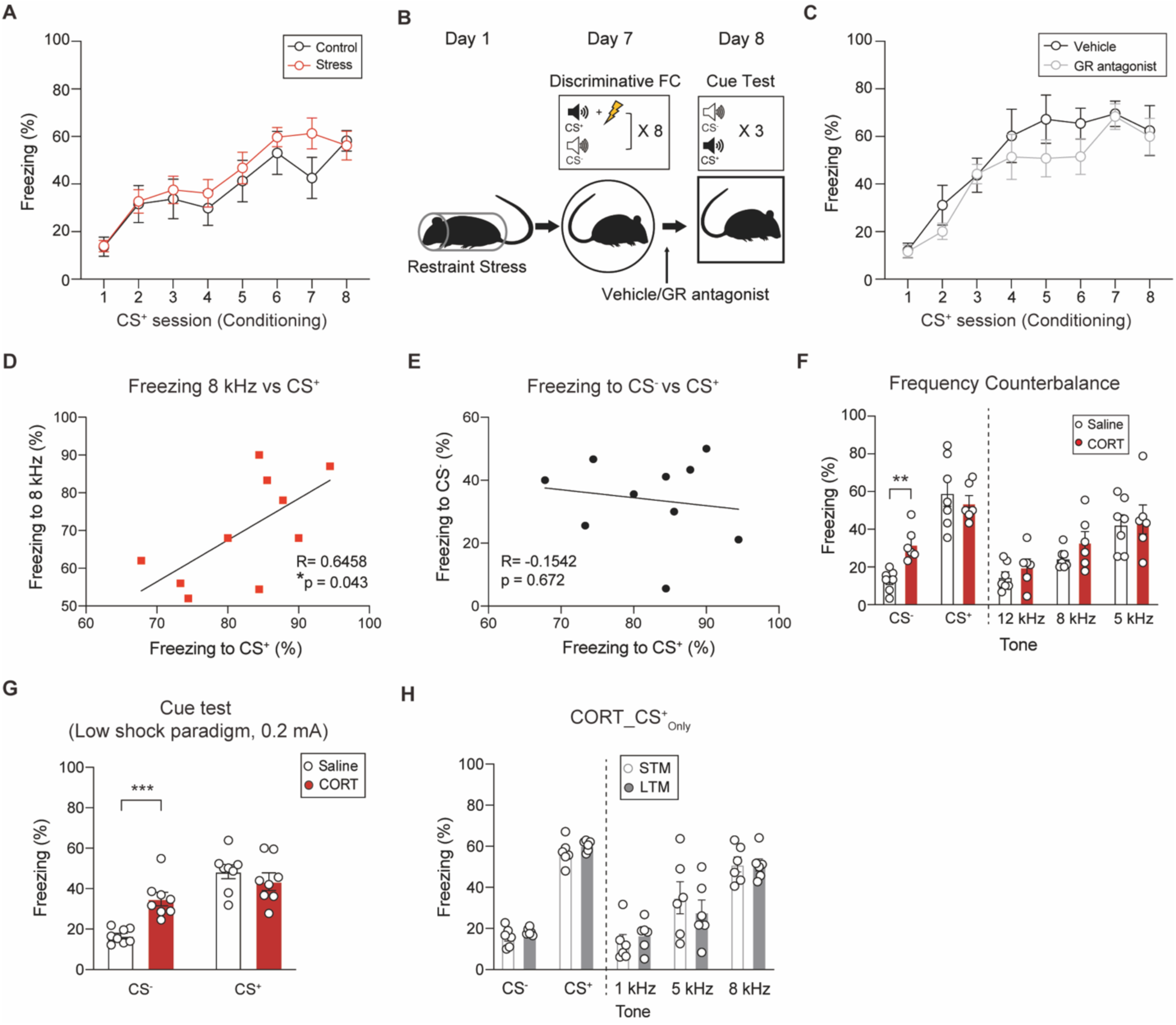
Glucocorticoid receptor-dependent elevation of fear expression to CS^-^. Related to **Figure 1**. **(A)** Freezing profiles to consecutive presentations of CS^+^ during discriminative FC between stressed and control mice (Control, n = 8; Stress, n =13 mice; Two-way Repeated Measure (RM) ANOVA, F = 0.6886, *P* = 0.4170). **(B)** Schematic depicting administration of a glucocorticoid receptor (GR) antagonist (mifepristone, 10 mg/kg) or vehicle in stressed mice and their behavioral tests. **(C)** Freezing profiles to consecutive presentations of CS^+^ during discriminative FC between mifepristone- and vehicle-injected stressed mice (Vehicle, n = 7; Mifepristone, n = 8 mice; Two-way RM ANOVA, F = 1.646, *P* = 0.2204). **(D - E)** Correlations of fear retrieval to CS^+^ and 8 kHz tones (D) or to CS^+^ and CS^-^ (E) in CORT-injected mice. **(F)** Fear retrieval in counter-balanced exposure to tones between saline- and CORT-injected mice (Saline, n = 7; CORT, n = 6 mice; Welch’s t test, \*\**P* = 0.0017). **(G)** Fear retrieval in mice underwent the low shock paradigm (0.2 mA) to tones between saline- and CORT-injected group (left, Saline, n = 8; CORT, n = 8; Welch’s *t*-test, \*\*\**P* = 0.0006). **(H)** Comparison of freezing levels to various tones at the STM and LTM time points in CORT_CS^+^_only_ group (left, n = 6 mice). Data are shown as mean ± SEM.

**Figure 1 - figure supplement 2.**
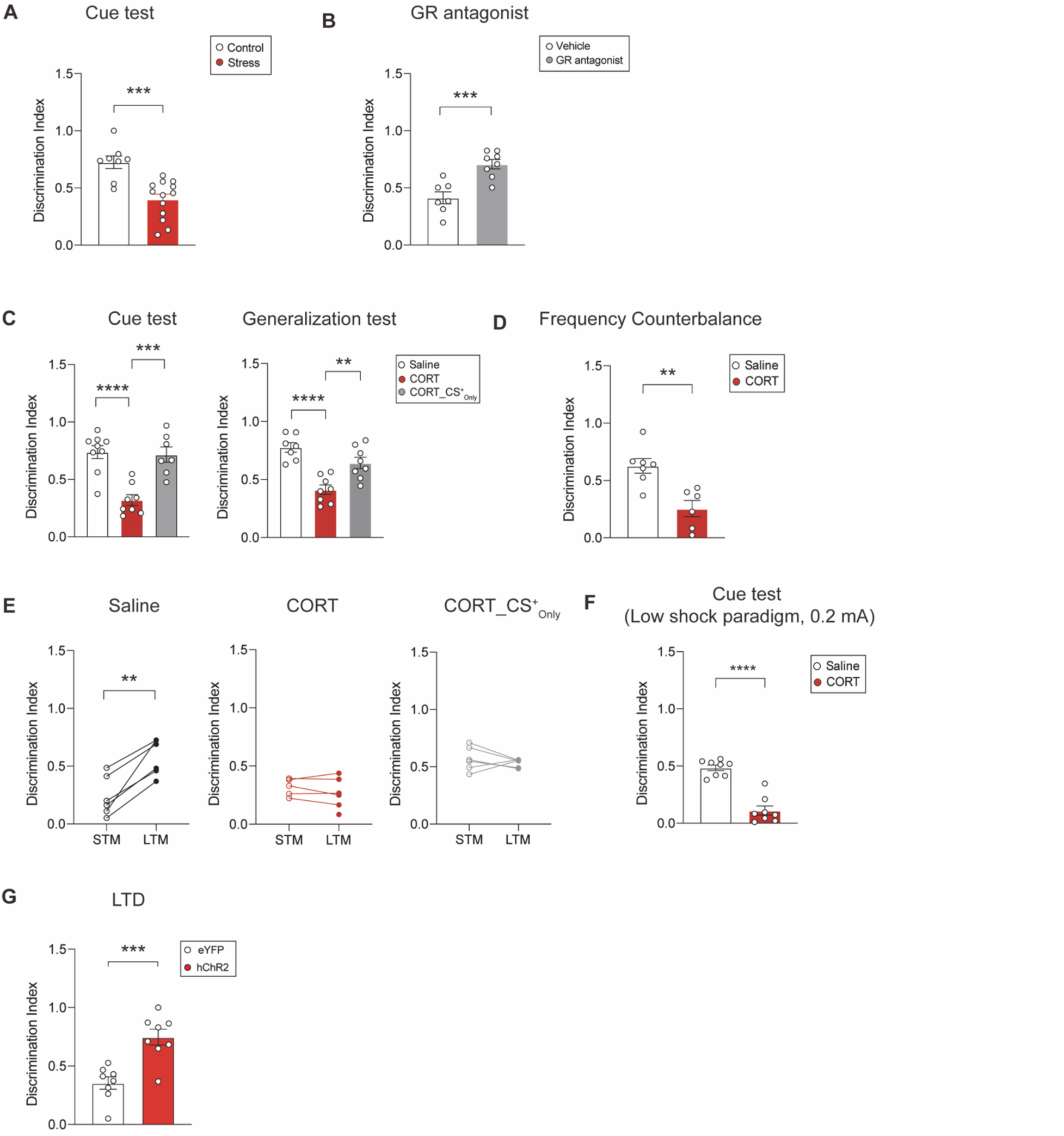
Comparison of Discrimination Index. Related to Figure 1. **(A)** Effects of stress exposure on discrimination index ((CS^+^ freezing – CS^-^ freezing) / (CS^+^ freezing + CS^-^ freezing)) during fear retrieval (Control, n = 8; Stress, n =13 mice; Welch’s *t*-test, \*\*\**P* = 0.0004). **(B)** Effects of GR blockage on discrimination index that stressed animals displayed during fear retrieval (Vehicle, n = 7; GR antagonist, n = 8 mice; Welch’s *t*-test, \*\*\**P* = 0.0009). **(C)** Comparison of discrimination index in the cue test (left, Saline, n = 9; CORT, n = 8; CORT_CS^+^_only_, n =7; One-way ANOVA with Tukey’s *posthoc* test, \*\*\**P* = 0.0003, \*\*\*\**P* < 0.0001) and the generalization test (right, Saline, n = 7; CORT, n = 8; CORT_CS^+^_only_, n =8; One-way ANOVA with Tukey’s *posthoc* test, \*\**P* = 0.0031, \*\*\*\**P* < 0.0001) from mice that underwent either discriminative FC or CS^+^_only_ training and then received CORT. **(D)** Comparison of discrimination index in counter-balanced exposure to tones (Saline, n = 7; CORT, n = 6 mice; Welch’s t test, \*\**P* = 0.0026). **(E)** Comparison of discrimination index at the STM and LTM time points in saline-injected (left, n = 6 mice; Paired *t*-test, \*\**P* = 0.002), CORT-injected (middle, n = 6 mice) and CORT_CS^+^_only_ groups (right, n = 6 mice). **(F)** Comparison of discrimination index in mice underwent the low shock paradigm (0.2 mA) (Saline, n = 8; CORT, n = 8; Welch’s *t*-test, \*\*\*\**P* < 0.0001). **(G)** Effect of LTD induction on discrimination index (right, eYFP, n = 8; hChR2, n = 8 mice; Welch’s *t*-test, \*\*\**P* = 0.0005). Data are shown as mean ± SEM.

**Figure 1 – figure supplement 3.**
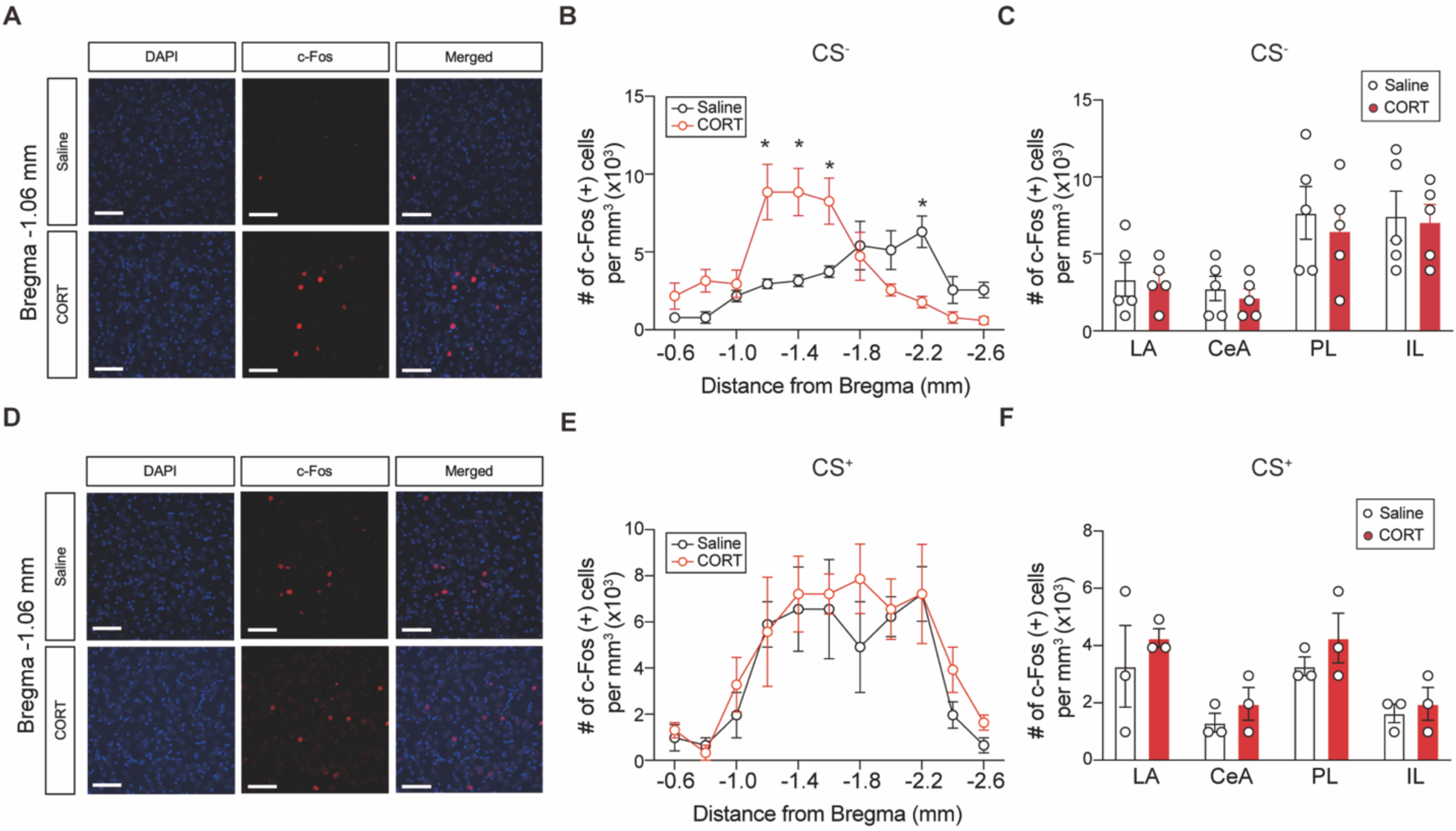
The c-Fos expression profiles in response to the CS^-^ and CS^+^. Related to Figure 1. **(A)** Representative images of DAPI staining and c-Fos-expressing cells in the aBA (bregma - 1.06 mm) after exposure to CS^-^. Scale bars, 40 μm **(B)** Topological comparison of c-Fos expression after CS^-^ exposure along the anterior to posterior axis of the BA (Saline, n = 5; CORT, n = 5 mice; Welch’s *t*-test, \**P* < 0.05). **(C)** Comparison of c-Fos-expressing cells after CS^-^ exposure in the LA, the central amygdala (CeA), the prelimbic cortex (PL) and the infralimbic cortex (IL)(Saline, n = 5; CORT, n = 5 mice). **(D)** Representative images of DAPI staining and c-Fos-expressing cells in the aBA (bregma -1.06 mm) after exposure to CS^+^. Scale bars, 40 μm. **(E)** Topological comparison of c-Fos expression after CS^+^ exposure along the anterior to posterior axis of the BA (Saline, n = 3; CORT, n =3 mice). **(F)** Comparison of c-Fos expression after CS^+^ exposure in various brain regions (Saline, n = 3; CORT, n =3 mice). Data are shown as mean ± SEM.

**Figure 1 - figure supplement 4.**
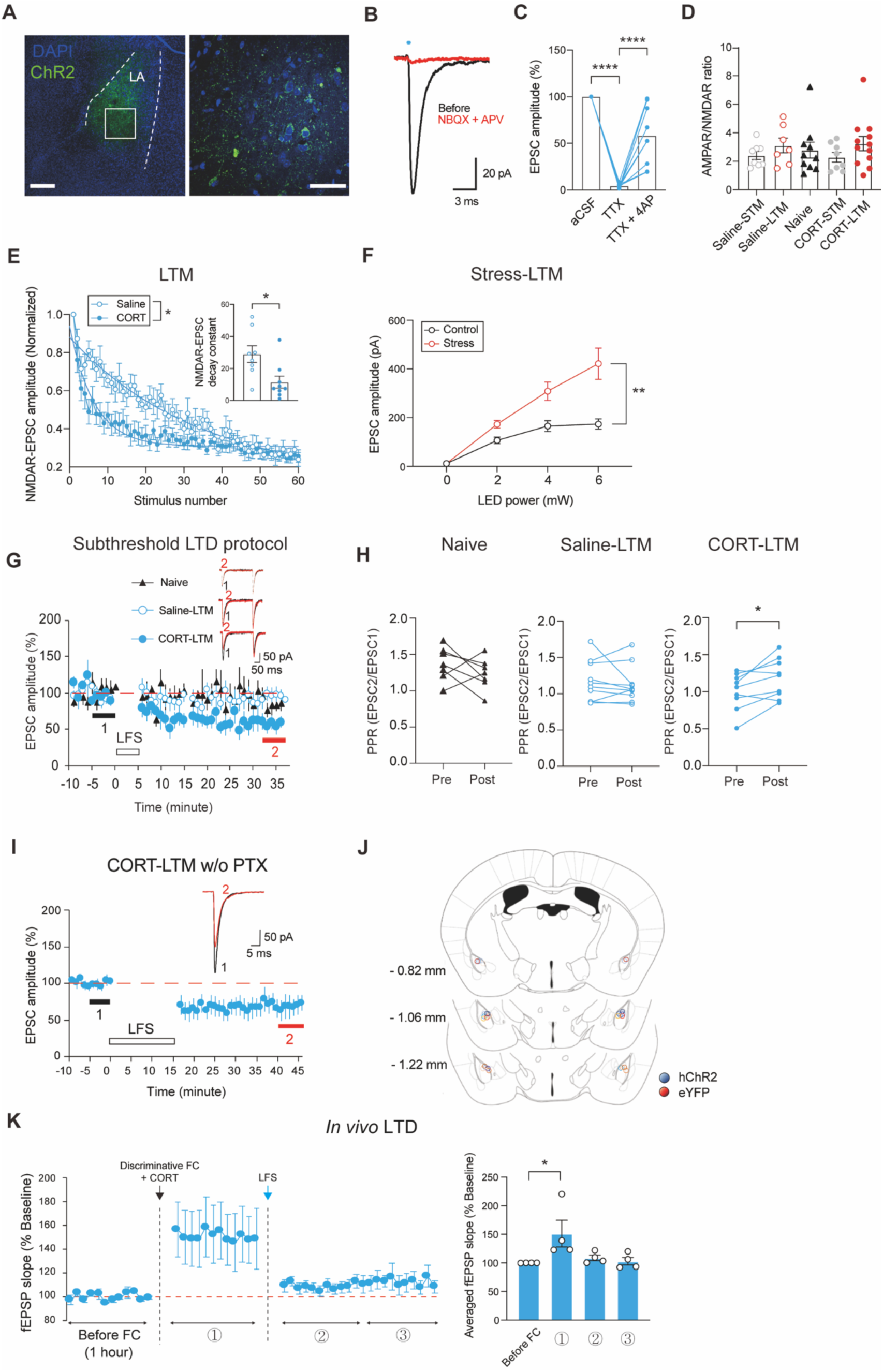
Presynaptic potentiation of the LA-to-aBA pathway associated with an increase in fear retrieval to CS^-^. Related to Figure 1. **(A)** Representative images for hChR2-eYFP expression. The LA is outlined with dotted lines (left) and the boxed area is magnified (right). Scale bars, 200 μm (left), 50 μm (right). **(B)** Representative EPSC traces elicited by optic stimulation of the LA-to-aBA pathway (in black), which was abolished by NBQX and APV (in red). **(C)** Monosynaptic excitatory transmission in the LA-to-aBA pathway which was blocked by Tetrodotoxin (TTX) and then partially restored by 4-aminopyridine (4-AP) (n = 9 cells; One-way RM ANOVA with Tukey’s *posthoc* test, \*\*\*\**P* < 0.0001). **(D)** Ratios of AMPAR/NMDAR-mediated EPSCs in each time point and condition (Naive, n = 10; Saline-STM, n = 8; Saline-LTM, n = 7; CORT-STM, n = 8; CORT-LTM, n = 12 cells). **(E)** Progressive blockade of NMDAR-EPSCs by MK801 between saline- and CORT-injected mice at the LTM time point (Saline, n = 8; CORT, n = 9 cells; Two-way RM ANOVA, F = 4.825, \**P* = 0.0442). Inserts: decay time constants of NMDAR-EPSCs (Saline, n = 8; CORT, n = 9 cells; Welch’s *t*-test, \**P* = 0.0173). **(F)** Input-output curves of EPSCs between stressed and unstressed control mice (Control, n = 9; Stress, n = 11 cells; Two-way RM ANOVA, F = 13.82, \*\**P* = 0.0016). **(G)** A subthreshold LTD protocol (low frequency stimulation, LFS) was applied onto the LA-to-aBA pathway of naive, saline- or CORT-injected mice at the LTM time point (Naive, n = 7; Saline-LTM, n = 10; CORT-LTM, n = 9 cells). Inserts: representative traces of EPSCs at the color-coded time points. **(H)** Paired pulse ratios (PPRs) before and after application of the subthreshold LTD protocol (Naive, n = 7; Saline-LTM, n = 10; CORT-LTM, n = 9 cells; Paired *t*-test, \**P* = 0.018). **(I)** *Ex vivo* validation of the *in vivo* LTD protocol. To recapitulate *in vivo* circumstances, low-frequency stimulation (LFS) was extended to 15 minutes in the absence of picrotoxin (CORT-LTM w/o PTX, n = 6 cells; values from 1 and 2 periods, Paired *t*-test, \**P* = 0.037). **(J)** Schematic depicting implantation sites of optical fibers within the LA. **(K)** fEPSP slopes before and after discriminative FC with CORT injection were measured witht subsequent LFS protocol (left, n = 4 mice). Averaged fEPSP slopes for each recording session (right, n = 4 mice; One-way repeated measures (RM) ANOVA with Tukey’s *posthoc* test, *\*P* = 0.0262). Data are shown as mean ± SEM.

**Figure 2 – figure supplement 1.**
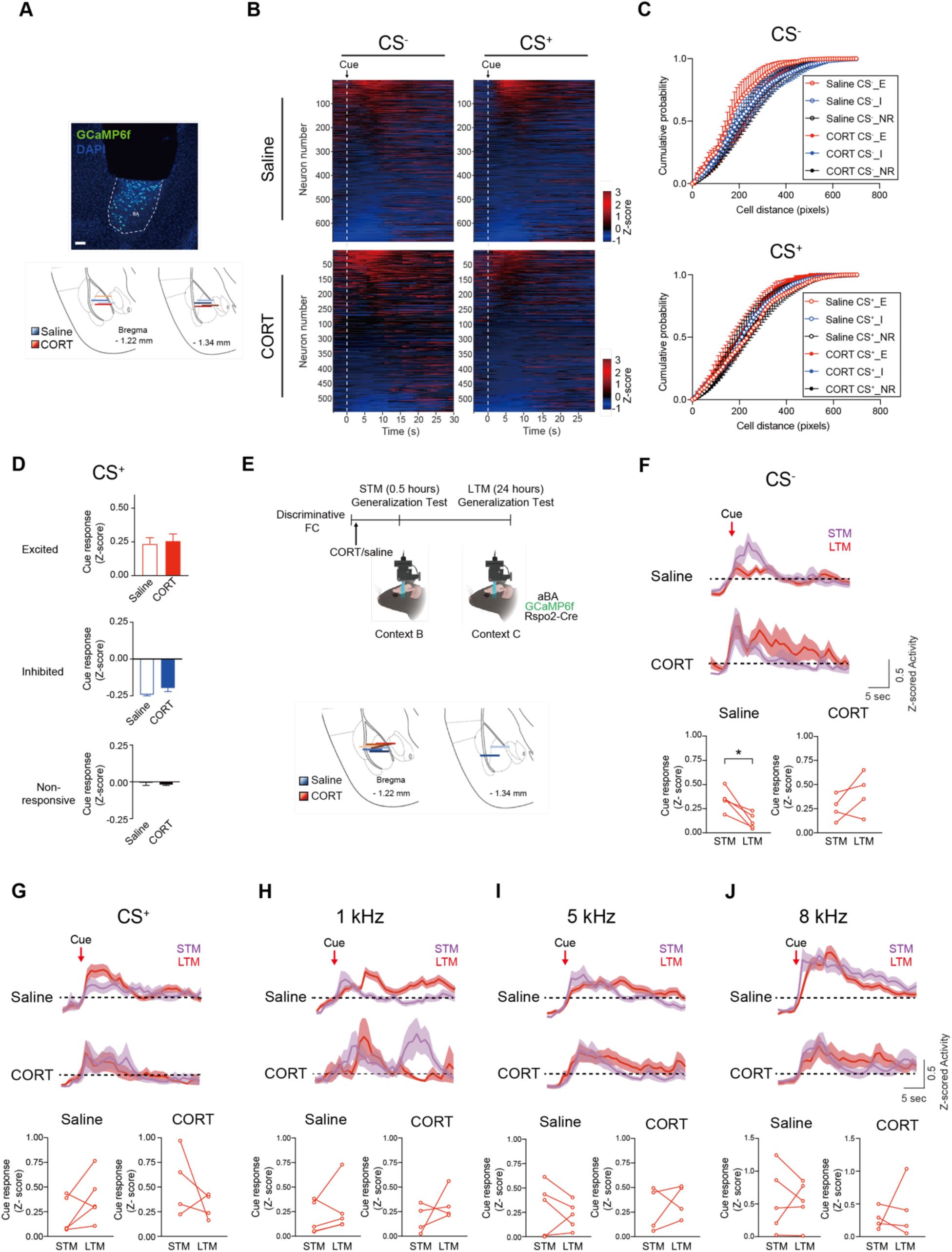
Cellular activity of Rspo2-expressing neurons upon exposure to cues. Related to Figure 2. **(A)** A representative image of a GRIN (Gradient index) lens implanted atop the BA expressing GCaMP6f. Scale bar, 100 μm (top). Schematic depicting positions of the bottoms of GRIN lenses in mice that received either saline or CORT (bottom). **(B)** Heatmaps of mean activity of aBA^Rspo2(+)^ neurons to CS^-^ or CS^+^ in saline- (left, n = 677) and CORT-(right, n = 543 cells) injected mice. Time points for cue exposure are indicated in arrows and dotted lines. Neurons are sorted according to their mean z-scored activity for 10 seconds after each stimulus onset. **(C)** The pairwise distance within each neural cluster to CS^-^ (top) or CS^+^ (bottom) in saline- and CORT-injected mice (Saline, n = 4; CORT, n = 4 mice). **(D)** Comparison of mean CS^+^-induced z-scored activity of each neural cluster between saline- and CORT-injected mice (top, Saline CS^+^_E, n = 120 *vs.* CORT CS^+^_E, n = 82; middle, Saline CS^+^_I, n = 183 *vs.* CORT CS^+^_I, n = 95; bottom, Saline CS^+^_NR, n = 374 *vs.* CORT CS^+^_NR, n = 366 cells). **(E)** An experimental timeline and schematic for *in vivo* Ca^2+^ imaging at the STM and LTM time points from aBA^Rspo2(+)^ neurons expressing GCaMP6f (top). Schematic depicting positions of the bottoms of GRIN lenses in mice that received either saline or CORT (bottom). **(F)** Averaged activity (z-scored) traces of excited-aBA^Rspo2(+)^ clusters to CS^-^ in the STM (purple) and LTM (red) points (top). Comparison of mean activity of excited clusters between two time points in saline- and CORT-injected mice (bottom, Saline n = 5; CORT, n = 4 mice; Paired *t*-test, \**P* = 0.0183). **(G-J)** Averaged activity (z-scored) traces of excited-aBA^Rspo2(+)^ clusters to CS^+^ (G), 1 Hz (H), 5 Hz (I) or 8 Hz tones (J) in the STM (purple) and LTM (red) sessions (top). Comparison of mean activity of excited clusters between two time points in saline- and CORT-injected mice (bottom, Saline n = 5; CORT, n = 4 mice). Data are shown as mean ± SEM.

**Figure 2 – figure supplement 2.**
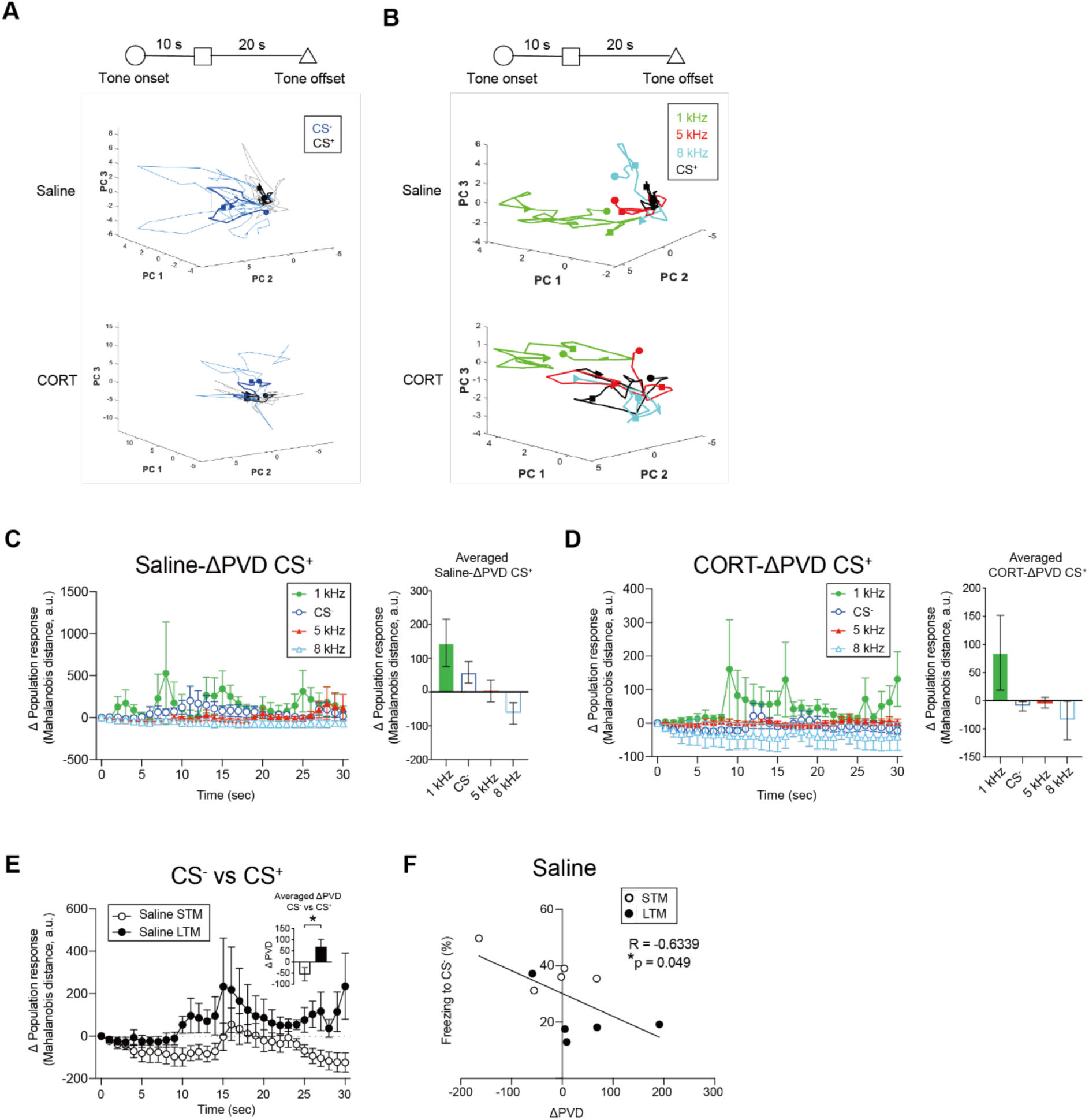
Population activity of Rspo2-expressing neurons upon exposure to cues. Related to Figure 2. **(A)** Representative neural trajectories of aBA^Rspo2(+)^ neurons to CS^-^ (blue) or CS^+^ (black) shown in lower dimensional PC spaces. The time points for activity measurement are designated by the symbols shown in the timeline (top). The light color indicates neural trajectory for individual trials and the dark color indicates the averaged traces in saline- (middle) and CORT- (bottom) injected mice. **(B)** Representative neural trajectories of aBA^Rspo2(+)^ neurons to tones of various frequencies (1 kHz, green; 5 kHz, red; 8 kHz, cyan; CS^+^, black) shown in low dimensional PC spaces in saline- (middle) and CORT- (bottom) injected mice. The time points for activity measurement are designated by the symbols shown in the timeline (top). **(C)** Deviation of population vector distance (ΔPVD) between CS^+^ and other tones (left) and comparison of the averaged ΔPVD (30 sec) in saline-injected mice (right) (n = 4 mice). **(D)** ΔPVD between CS^+^ and other tones (left) and comparison of the averaged ΔPVD (30 sec) in CORT-injected mice (right) (n = 4 mice). **(E)** ΔPVD between CS^-^ and CS^+^ in the STM and LTM time points in saline-injected mice. Insert: comparison of averaged ΔPVD between STM and LTM points in saline-injected mice (n = 5 mice; Paired *t*-test, \**P* = 0.0377). **(F)** Inverse correlation between freezing magnitudes to CS^-^ measured at two time points and the corresponding ΔPVD. Data are shown as mean ± SEM.

**Figure 3 – figure supplement 1.**
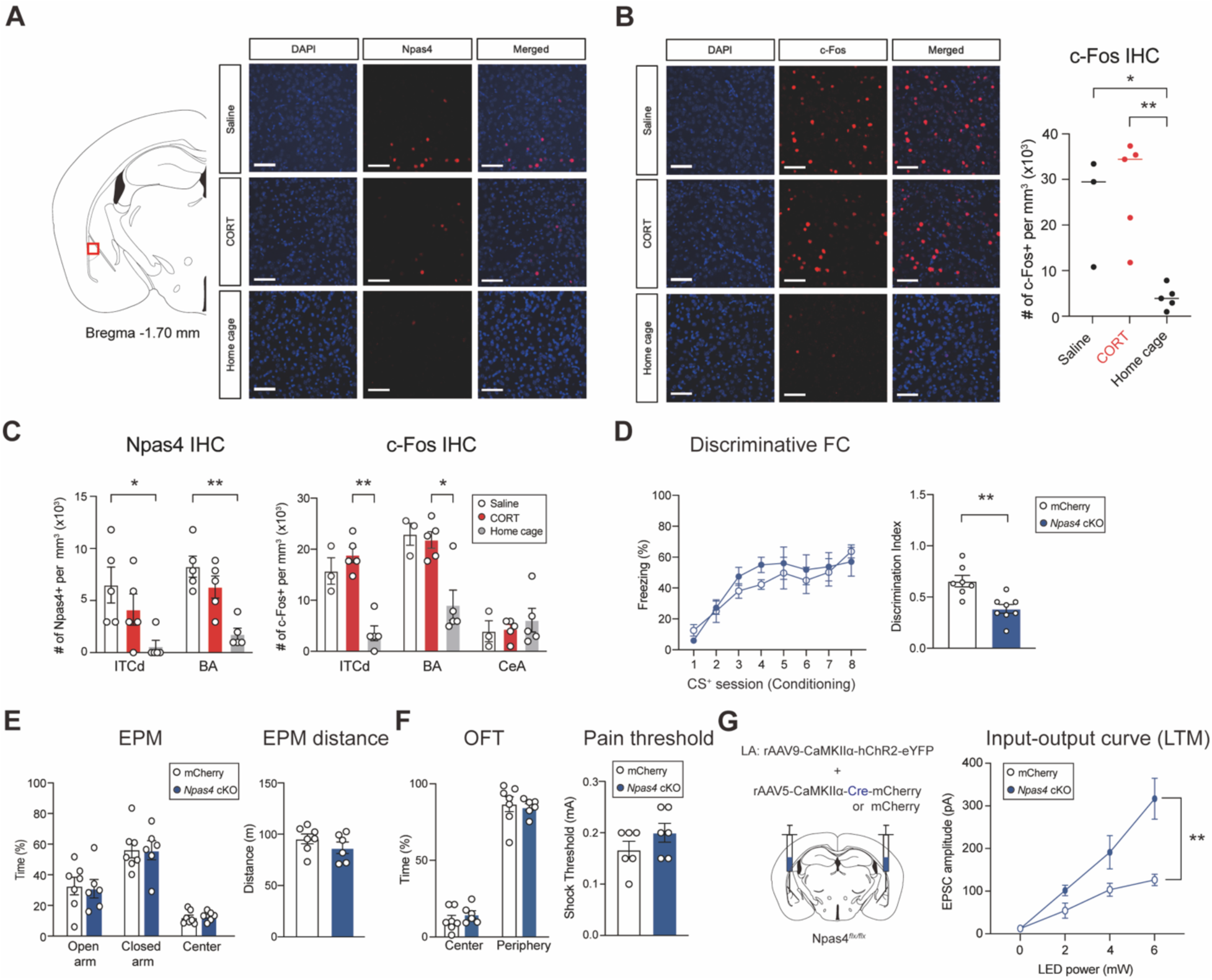
Expression profiles of IEGs and effect of Npas4 on behaviors. Related to Figure 3. **(A)** Representative images of Npas4-expressing cells in the LA after discriminative fear conditioning (FC). Scale bars, 40 μm. **(B)** Representative images of c-Fos expression in the LA. Scale bars, 40 μm (left). Comparison of c-Fos expression in the LA 90 minutes after discriminative FC in saline- and CORT-injected mice as well as home cage group (right, Saline, n. = 3; CORT, n = 5; Home cage, n = 5 mice; One-way ANOVA with Tukey’s *posthoc* test, \*\**P* = 0.0267, \*\**P* = 0.0045). **(C)** Comparison of Npas4 expression between saline-, CORT-injected and home cage groups 90 minutes after discriminative FC (left, Saline, n = 5; CORT, n = 5; Home cage, n = 5 mice; Kruskal-Wallis test with Dunn’s *posthoc* test, \**P* = 0.0244, \*\**P* = 0.0094) and c-Fos expression (right, Saline, n = 3; CORT, n = 5; Home cage, n = 5 mice; Kruskal-Wallis test with Dunn’s *posthoc* test, \**P* = 0.0345, \*\**P* = 0.0095). **(D)** Freezing profiles to consecutive presentations of CS^+^ during discriminative FC between *Npas4* cKO and mCherry control mice (left, *Npas4* cKO, n = 8; mCherry, n = 7 mice). Effect of Npas4 cKO in discrimination index during fear retrieval (right, *Npas4* cKO, n = 8; mCherry, n = 7 mice; Welch’s *t*-test, \*\**P* = 0.0025). **(E)** Time duration spent at each sector (left) and traveled distance (right) in EPM test (*Npas4* cKO, n = 6; mCherry, n = 7 mice). **(F)** Time duration that animals spent in the center or periphery areas of OFT assay (left) and shock thresholds in a pain test (right, *Npas4* cKO, n = 6; mCherry, n = 7 mice). **(G)** Schematic for recording synaptic transmission of the LA-to-aBA pathway (left). Input-output curves of EPSCs between *Npas4* cKO and mCherry control mice (right, *Npas4* cKO, n = 8; mCherry, n = 8 cells, Two-way RM ANOVA, F = 13.58, \*\**P* = 0.0025). Data are shown as mean ± SEM.

**Figure 4 – figure supplement 1.**
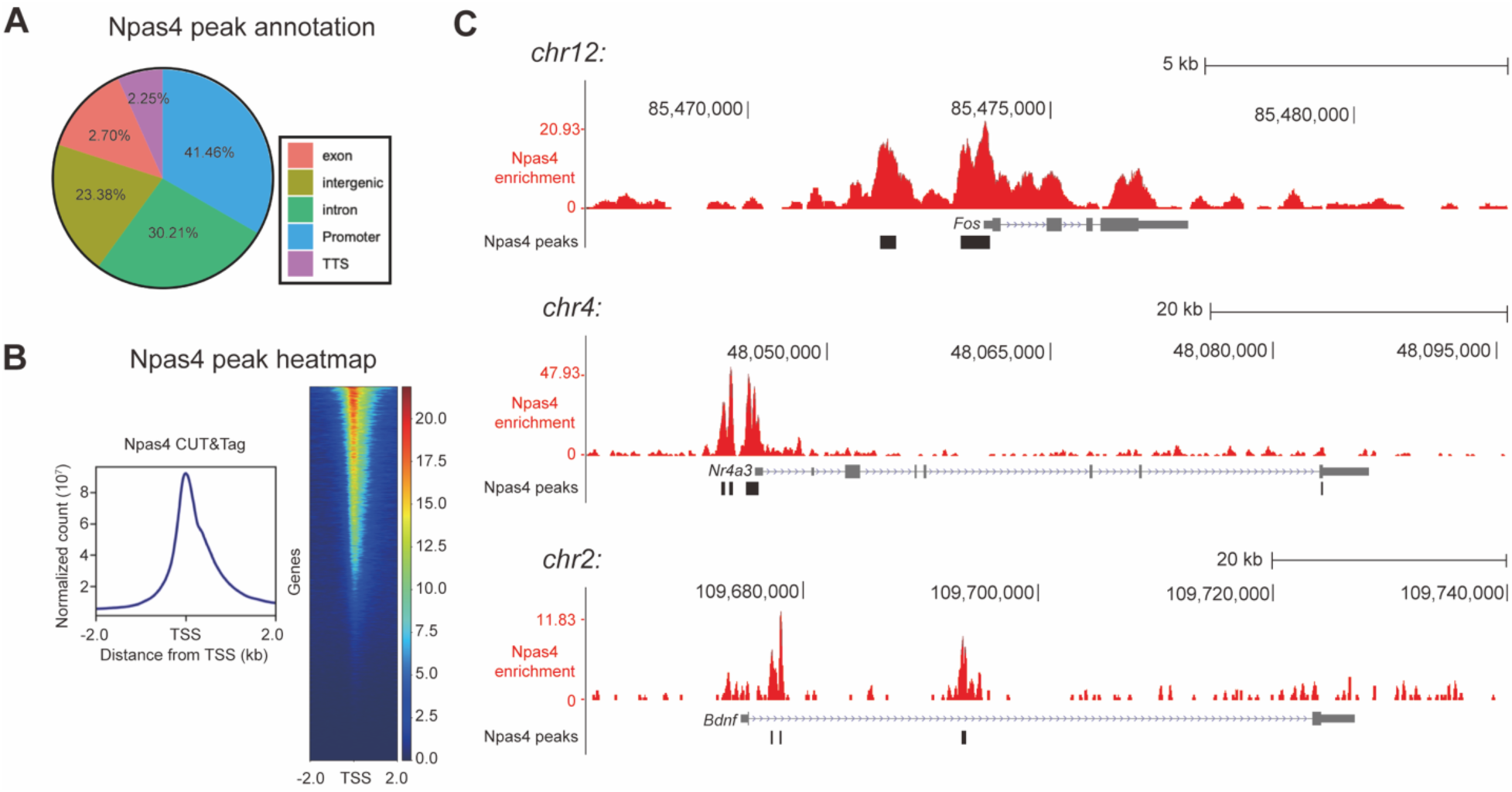
CUT&Tag identified Npas4-enriched regions in amygdala. Related to Figure 4. **(A)** Genomic annotation of Npas4 peaks identified by CUT&Tag analysis in accordance with RefSeq genes. **(B)** An aggregate plot showing spike-in normalized counts across all Npas4 peaks (left) and a heatmap over 2 kb regions spanning the center of transcription start sites (TSS) (right). **(C)** Npas4 binding loci surrounding the *c-Fos*, (top), the *Nr4a3* (middle) or, the *BDNF* (Brain-derived neurotrophic factor, bottom) promoters. Y-axis indicates normalized reads (10^7^). Data are shown as mean ± SEM.

**Figure 4 – figure supplement 2.**
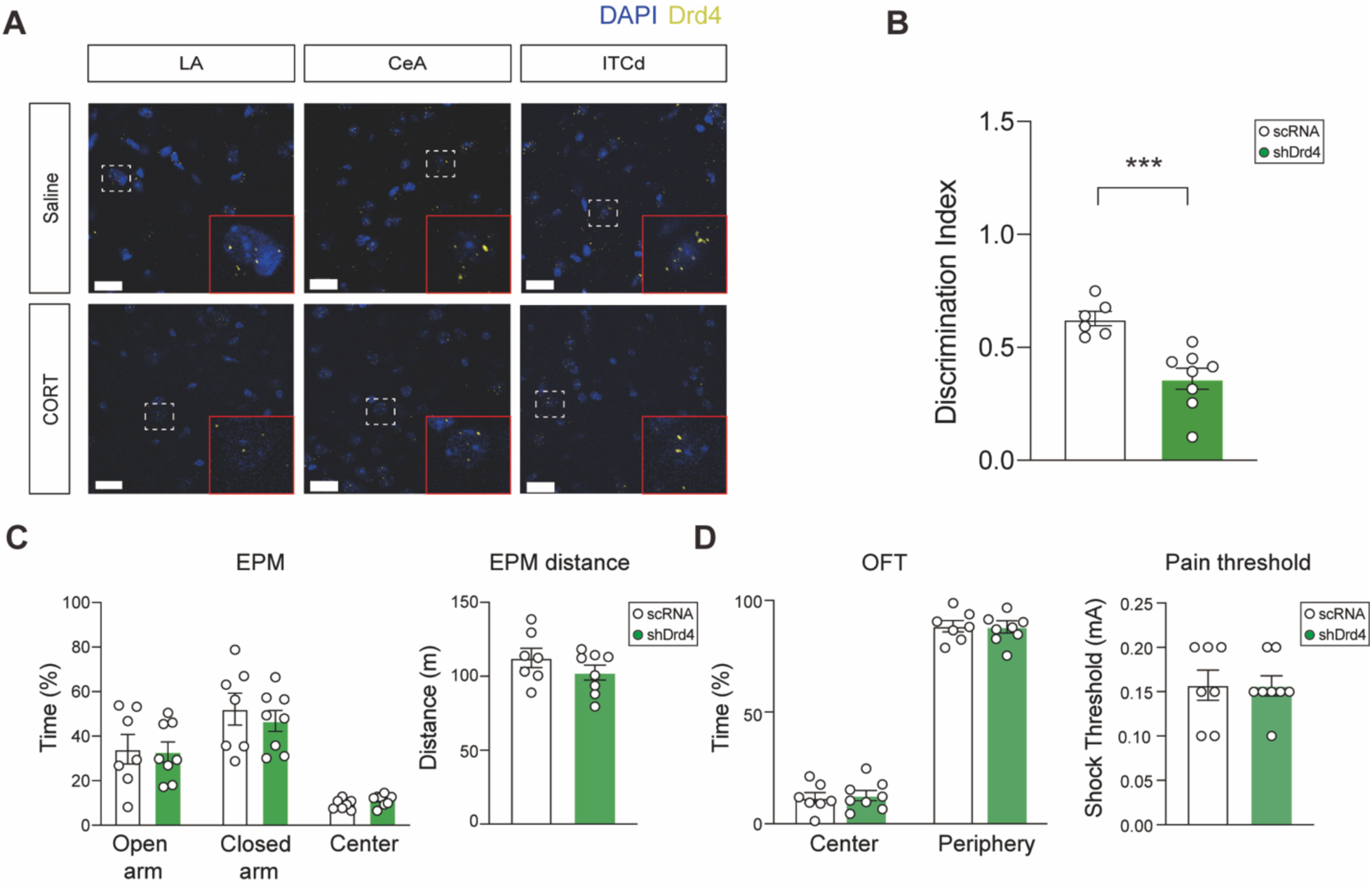
Effects of Drd4 signaling on behaviors. Related to Figure 4. **(A)** Representative images for Drd4 puncta in amygdala sub-regions of saline- and CORT-injected mice revealed by immunohistochemistry. Areas outlined with white dotted lines are amplified into those with red solid lines. Scale bars, 20 μm. **(B)** Comparison of discrimination index between shD4R- and scRNA-infused mice ((shDrd4, n = 8; scRNA, n = 6; Welch’s *t*-test, \*\*\**P* = 0.0006). **(C)** Time duration spent at each sector (left) and traveled distance (right) in EPM test between shD4R- and scRNA-infused mice (shDrd4, n = 8; scRNA, n = 7 mice). **(D)** Time duration that animals spent in the center or periphery areas of OFT assay (left) and shock thresholds in a pain test (right)(shDrd4, n = 8; scRNA, n = 7 mice). Data are shown as mean ± SEM.

**Figure 5 – figure supplement 1.**
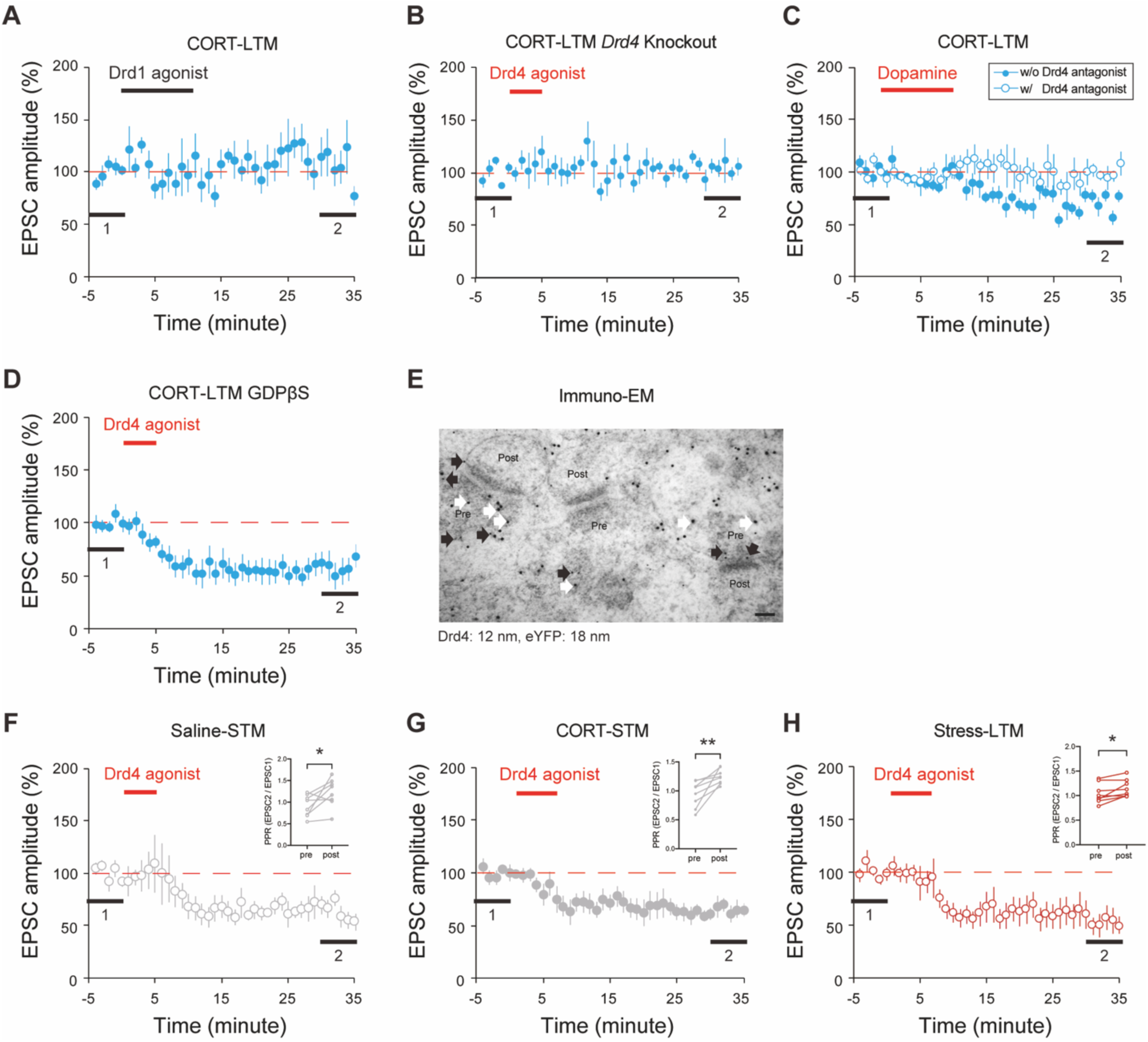
Effects of Drd4 signaling on synaptic transmission. Related to Figure 5. **(A)** Effects of Drd1 agonist (SKF-38393) on synaptic transmission in CORT-LTM group (n = 8 cells; 1 *vs* 2, Paired *t*-test, *P* = 0.6915). **(B)** Effects of Drd4 agonist (PD-168,077) on synaptic transmission of *Drd4* KO mice in CORT- LTM group (n = 7 cells; 1 *vs* 2, Paired *t*-test, *P* = 0.4502). **(C)** Effects of dopamine on synaptic transmission in presence or absence of Drd4 antagonist (L- 745,870) in CORT-LTM group (without L-745,870: n = 9; 1 *vs* 2, Paired *t*-test, \*\*\*\**P* < 0.0001 *vs.* with L-745,870: n = 6 cells; 1 *vs* 2, Paired *t*-test, *P* = 0.9411). **(D)** PD-168,077-induced LTD indifferent to postsynaptic GCPR signaling (n = 7 cells; 1 *vs* 2, Paired *t*-test, \*\**P* = 0.0087). **(E)** Immuno-electron microscopic (EM) images showing the abundant distribution of Drd4 in the presynaptic areas marked with eYFP. Black and white arrows denote Drd4-immunogold (12 nm) and eYFP-immunogold particles (18 nm), respectively. Scale bars, 100 nm. **(F - G)** Effects PD-168,077 on synaptic transmission in Saline-STM (F) (n = 8 cells; 1 *vs* 2, Paired *t*-test, \*\*\**P* = 0.0003) and CORT-STM (G) groups (n = 9 cells; 1 *vs* 2, Paired *t*-test, \*\*\**P* = 0.0002). Inserts: PPRs measured before and after perfusion (Saline-STM, n = 8, Paired *t*-test, \**P* = 0.0461; CORT-STM, n = 9 cells, Paired *t*-test, \*\**P* =0.0077). **(H)** Effects of PD-168,077 on synaptic transmission in stress-LTM group (n = 8 cells; 1 *vs* 2, Paired *t*-test, \*\*\**P* = 0.0004). Insert: PPRs measured before and after the perfusion (n = 8 cells; Paired *t*- test, \**P* = 0.0335). Data are shown as mean ± SEM.

**Figure 6 – figure supplement 1.**
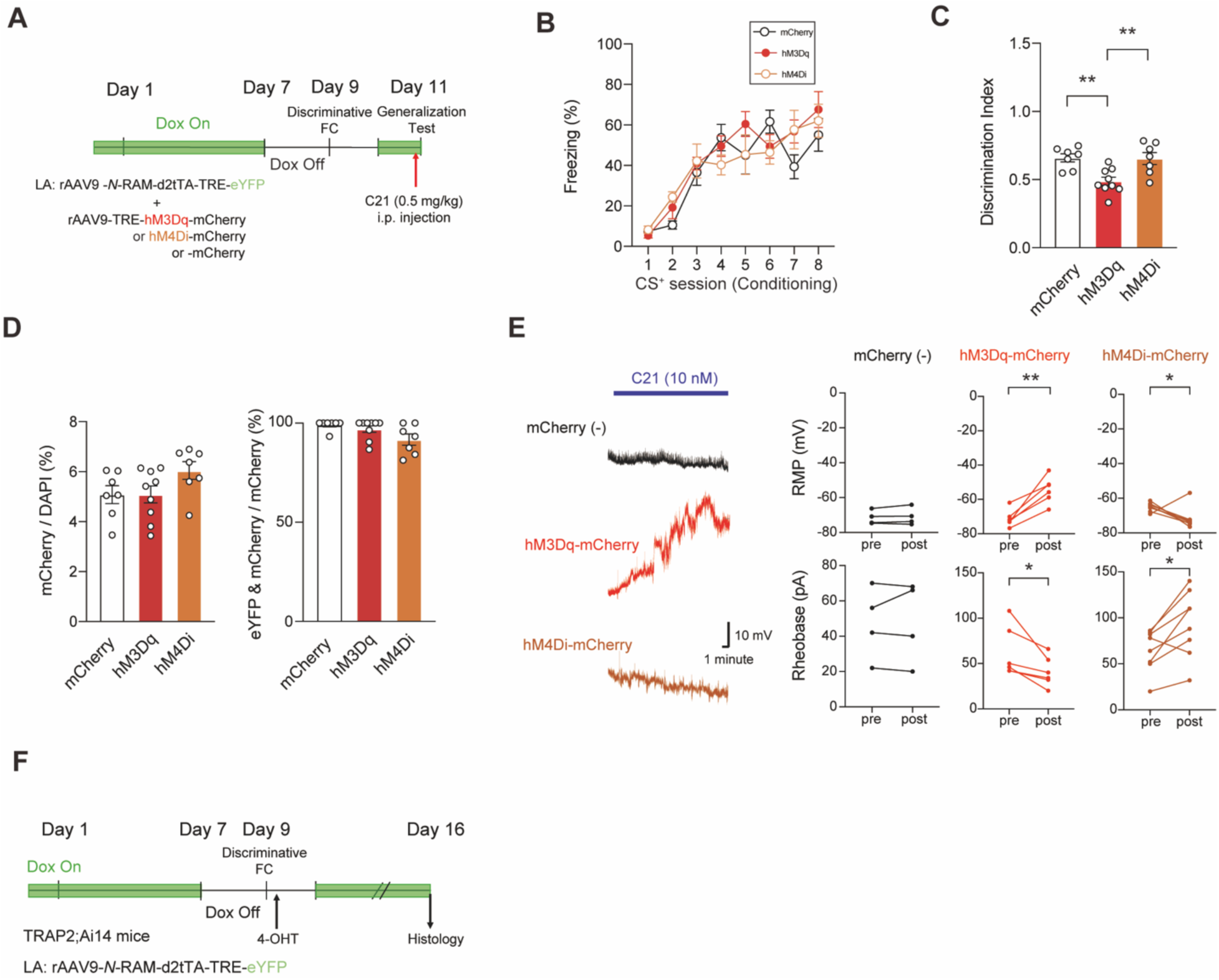
Functionality validation of *N*-RAM system. Related to Figure 6. **(A)** An experimental timeline for activity manipulation of *N*-RAM-labelled cells in the LA. **(B)** Freezing profiles during fear conditioning in mice expressing mCherry-, hM3Dq- or hM4Di in the LA (mCherry, n = 7; hM3Dq, n = 9; hM4Di, n = 7 mice). **(C)** Comparison of discrimination index during fear retrieval among groups (mCherry, n = 7; hM3Dq, n = 9; hM4Di, n =7 mice; One-way ANOVA with Tukey’s *posthoc* test, \*\**P* < 0.01). **(D)** Efficacy of viral labeling among groups. Infection ratios (numbers of mCherry-labelled cells relative to numbers of DAPI-stained cells) for each group (left), and specificity of *N-*RAM-mediated labeling (right, mCherry, n = 7; hM3Dq, n = 9; hM4Di, n = 7 mice). **(E)** Representative traces for resting membrane potentials (RMPs) with C21 treatment (left). RMPs (top) and rheobases (bottom) between before and after C21 treatment for each group (right, mCherry-negative, n = 4; hM3Dq-mCherry, n = 6; hM4Di-mCherry, n = 8 cells; Paired *t*-test, \**P* < 0.05, \*\**P* = 0.0026). **(F)** An experimental timeline for simultaneous labeling of Npas4- and c-Fos-expressing cells in the LA. Data are shown as mean ± SEM.

## Notes

### Competing Interest Statement

The authors have declared no competing interest.

